# CHRM1 is a Druggable Melanoma Target Whose Endogenous Activity is Determined by Inherited Genetic Variation in DOPA Production

**DOI:** 10.1101/2021.03.03.433757

**Authors:** Miriam Doepner, Christopher A. Natale, In Young Lee, Swati Venkat, Sung Hoon Kim, John A. Katzenellenbogen, Benita S. Katzenellenbogen, Michael E. Feigin, Todd W. Ridky

## Abstract

Melanoma risk is 30 times higher in people with lightly pigmented skin compared to those with darkly pigmented skin. Here we show that this difference results from more than melanin pigment and its ultraviolet radiation (UVR) shielding effect. Using primary human melanocytes representing the full human skin pigment continuum and several preclinical melanoma models, we show that cell-intrinsic differences between dark and light melanocytes regulate melanocyte proliferative capacity, overall cellular differentiation state, and susceptibility to malignant transformation, <u>independently of melanin and UV exposure.</u> We determined that these differences result from dihydroxyphenylalanine (DOPA), a melanin precursor synthesized at higher levels in melanocytes from dark skin. Although DOPA was not previously known to have specific signaling activity, we used both high throughput pharmacologic and genetic *in vivo* CRISPR screens to determine that DOPA limits melanocyte and melanoma cell proliferation by directly inhibiting the muscarinic acetylcholine receptor M1 (CHRM1), a G Protein-Coupled Receptor (GPCR) not previously known to bind DOPA, nor to affect melanoma pathobiology. Pharmacologic CHRM1 antagonism in melanoma leads to depletion of c-Myc and FOXM1, both of which are proliferation drivers associated with aggressive melanoma. In preclinical mouse melanoma models using both immune deficient and syngeneic immune competent mice, pharmacologic inhibition of CHRM1 or FOXM1 inhibited tumor growth. CHRM1 and FOXM1 may be new therapeutic targets for melanoma.

## Introduction

Melanoma is the most lethal form of skin cancer on a per case basis. Despite advances in modern immune and targeted therapies, most patients with metastatic melanoma still die from their disease and new treatment approaches are needed.^1,2^ Clues to new therapeutic approaches may lie in understanding the mechanisms by which melanoma differentially affects different populations of people. Here we consider why the lifetime risk for cutaneous melanoma is substantially higher for people with lightly pigmented skin compared to those with darkly pigmented skin, even when they live in the same geographic region and are thereby exposed to similar amounts of UVR.^3^

Melanoma develops from melanocytes (MCs), which normally reside in the basal layer of skin and hair follicles where they produce melanin pigment, the primary determinant of skin and hair color. Melanogenesis is a complex, multi-step process that begins with the non-essential amino acid L-tyrosine and results in the production of mostly insoluble eumelanin (brown-black) or pheomelanin (red-yellow) polymers.^4–6^ Variation in the eumelanin to pheomelanin ratio creates the natural diversity in human skin pigmentation. These baseline pigmentary differences result from numerous single nucleotide polymorphisms (SNPs) in at least 200 genes involved in melanin synthesis.^7^ Eumelanin acts as a physical photoprotective filter against DNA damaging solar ultraviolet radiation (UVR) and thereby protects skin cells from deleterious mutations that may lead to malignant transformation.^8^ While melanin’s UVR shielding effect undoubtedly accounts for some of the differences in lifetime melanoma risk across the diverse human pigment continuum, highly pigmented skin provides a sun protective factor (SPF) of only 2-3 versus lightly pigmented skin, which seems insufficient to completely explain the large 30-fold difference in skin cancer incidence between people with lightly pigmented vs darkly pigmented skin.^9,10^ Furthermore, a UVR shielding effect does not fully explain decades of epidemiologic data suggesting that there are UV-independent determinants of melanoma risk that also correlate with skin pigment type. Melanomas arising in completely sun-protected areas, such as anorectal melanoma, are up to 13 times more common in people with lightly vs highly pigmented skin.^11,12^ There is also an intriguing observation involving skin cancer in people from Africa with albinism. While affected individuals have epidermal MCs, they do not make melanin, and therefore have white or extremely lightly pigmented skin and hair. They exhibit photosensitivity and an expected elevated incidence of keratinocyte-derived cancers, including basal cell and squamous cell carcinomas. However, they appear highly resistant to melanoma suggesting that while their MCs are visibly light, they may be functionally “dark” with regard to melanoma, and thereby similar to those with darkly pigmented skin in their population group with shared African ancestry.^13,14^ The mechanism(s) underlying these apparent UV-independent determinants of melanoma susceptibility were previously unknown.

Here, we show that endogenously produced dihydroxyphenylalanine (DOPA), a melanin synthesis intermediate, drives cellular differentiation in primary human MCs, which is associated with slower proliferation and resistance to the oncogenic effects of the major human melanoma oncoprotein BRAF(V600E). We show that these DOPA effects result from antagonism of the muscarinic acetylcholine receptor M1 (CHRM1), a G Protein-Coupled Receptor (GPCR) on melanocytes and melanoma cells. In preclinical *in vivo* melanoma models, pharmacologic CHRM1 antagonism inhibited melanoma growth. We show that inhibition of CHRM1 induced depletion of FOXM1, a transcription factor and cell cycle regulator associated with more aggressive cancer, and that a new class of FOXM1 specific antagonists also significantly inhibited melanoma growth *in vivo* and extended overall survival. Together these data suggest that CHRM1 and FOXM1 may be new druggable targets for melanoma and emphasize that differences in melanoma risk across the human skin pigment continuum are more complex than can be explained simply a physical UV shielding effect from melanin.

## Results

### Darkly pigmented MCs are less tumorigenic than light pigmented MCs

Under standard cell culture conditions without UVR, lightly pigmented early passage primary human MCs (LMC) proliferated 2-3 times faster than darkly pigmented MCs (DMC) **(Figure 1a)**. Melanocyte proliferative capacity is classically inversely correlated with MC cellular differentiation state^15–19^, which is primarily regulated by the activation of Gs-coupled G protein-coupled receptors (GPCRs).^20–23^ Gs signaling stimulates production of cyclic adenosine monophosphate (cAMP) via adenylate cyclase. In MCs, cAMP activates protein kinase A (PKA), which phosphorylates and activates the cAMP response element-binding protein (CREB), which ultimately promotes downstream synthesis of proteins involved in melanin production, such as tyrosinase (Tyr).^24,25^ We examined whether the expression of proteins within this classic GPCR pathway differed between LMCs and DMCs. DMCs contained more phosphorylated CREB (pCREB) and tyrosinase than LMCs **(Supplemental Figure 1a)**, suggesting that DMCs are more fully advanced along a cellular differentiation continuum that parallels the natural range of human skin pigment diversity. Consistent with this idea, DMCs expressed less of the stem cell marker and oncoprotein c-Myc **(Figure 1b, Supplemental Figure 1a)**.

**Figure 1:**
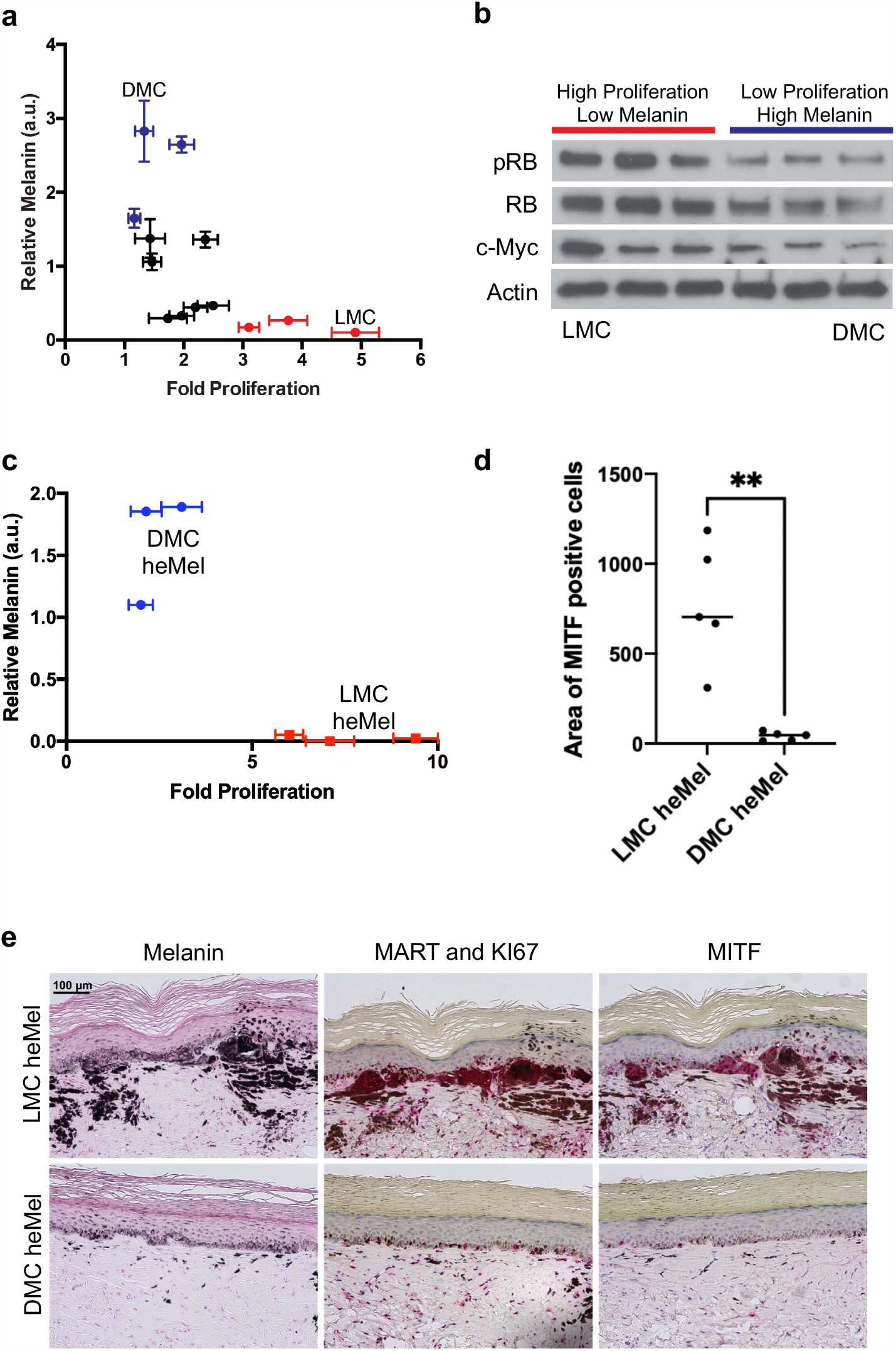
Cell-intrinsic differences render DMCs less tumorigenic than LMCs. (a) Scatter plot of 12 individual primary human melanocytes cultures shows melanin content at 450 nm wavelength vs. proliferation capacity. (b) Western blot of proliferation markers in lightly pigmented melanocytes (LMC) and darkly pigmented melanocytes (DMC) at baseline. (c) Scatter plot of transformed human engineered melanoma (heMel) shows melanin content at 450 nm wavelength vs. proliferation capacity. (d) Quantification of positive epidermal MITF staining area compared to total epidermal area in LMC and DMC heMel samples. P-value ** = 0.0082 analyzed via t-test. (e) Histologic characterization of representative orthotopic skin and resulting tumors, including melanocyte and proliferation markers MITF, Ki67/MART, and Fontana Masson (Melanin). Images taken at 20x magnification. Scale bar = 100 µm.

We hypothesized that these baseline differences in relative cellular differentiation state and proliferative capacity between DMCs and LMCs contribute to overall differential melanoma susceptibility. To test this *in vivo*, we used a genetically defined orthotopic human melanoma (heMel) model.^26,27^ Primary LMC and DMCs were engineered using lentiviruses to express mutant oncoproteins associated with spontaneous human melanoma including BRAF^V600E^, dominant-negative p53^R248W^, active CDK4^R24C^, and hTERT^26^. Equal expression of all oncoproteins was confirmed in darkly pigmented and lightly pigmented heMel cells **(Supplemental Figure 1c)**. The proliferation and differentiation differences between DMCs and LMCs observed in the untransduced parental cells remained after transduction of the oncoproteins. Darkly pigmented heMel cells proliferated over two times slower than lightly pigmented heMel cells and maintained their more differentiated phenotype **(Figure 1c, Supplemental Figure 1b)**, suggesting that cell intrinsic factors in DMCs may protect them from the oncogenic effects of common melanoma drivers. To test whether these *in vitro* differences translated to different melanoma phenotypes *in vivo*, lightly and darkly pigmented heMel cells were combined with normal primary human keratinocytes and incorporated into devitalized human dermis to establish 3-dimensional skin tissues in organotypic culture.^28^ We then grafted the engineered skin onto the orthotopic location on the backs of severe combined immunodeficient (SCID) mice. After 100 days, the grafts were harvested and analyzed histologically. Tissues with lightly pigmented heMel cells formed early melanomas, with large proliferative melanocytic nests, defined by MITF and MART staining, and with hallmark melanoma features, including early dermal invasion and upward Pagetoid scatter **(Figure 1d, e)**. In striking contrast, darkly pigmented heMel cells did not progress to melanoma, although the individual heMel cells were present in the tissues **(Figure 1d, e)**. These results show that DMCs resist BRAF-driven transformation, independent of UVR.

### DOPA inhibits MC proliferation and melanoma *in vitro* and *in vivo*

Although dark MCs contain more pigment than light MCs, eumelanin is a highly insoluble, large heterogeneous polymer without known signaling activity. Therefore, to begin defining the mechanism(s) responsible for reduced proliferation in DMCs, we first we looked to upstream intermediates in the melanin synthesis pathway. Melanin is synthesized via a complex multistep process involving serial oxidation and polymerization of tyrosine and is regulated by over 200 different genes **(Figure 2a)**.^29^ Tyrosine is first converted into L-dihydroxyphenylalanine (DOPA) via tyrosinase, and this is the rate limiting step in melanin synthesis.^23,30^ Consistent with the premise that tyrosinase activity increases in parallel with eumelanin content across the human pigment spectrum^6,31^, we detected approximately 300% more DOPA in cultures of primary human DMCs, as compared to LMCs **(Figure 2b)**.

**Figure 2:**
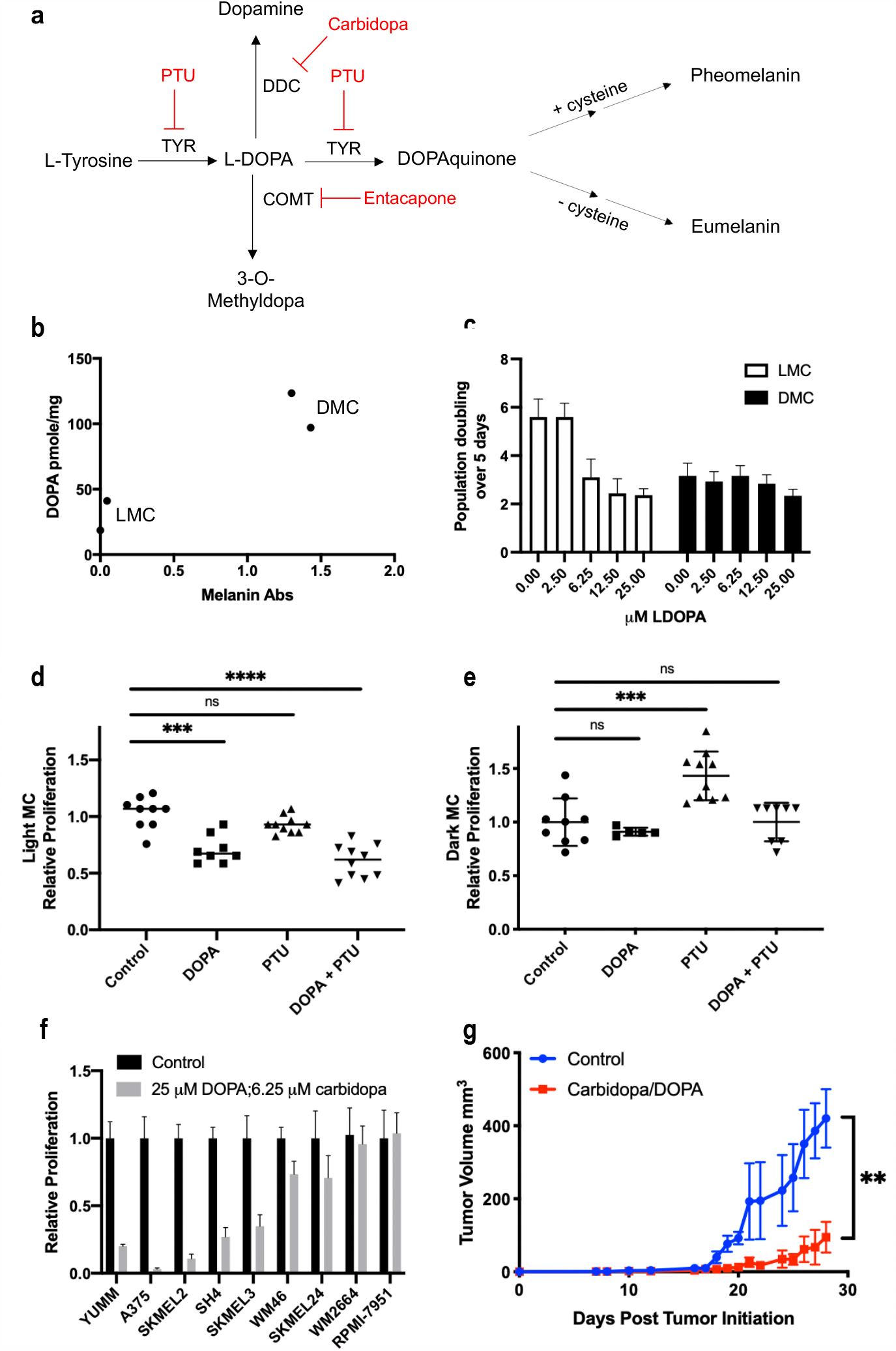
DOPA inhibits MC proliferation and melanoma *in vitro* and *in vivo*. (a) Schematic diagram depicting melanin synthesis. Pharmacologic inhibitors used in this paper are shown in **red**. (b) LC-MS quantitation of DOPA content in lightly pigmented melanocytes (LMC) and darkly pigmented melanocytes (DMC). (c) Dose curve of L-DOPA in representative LMC and DMC after 4 days L-DOPA treatment. (d) LMCs treated with either 25 µM L-DOPA, 75 µM phenylthiourea (PTU), or a combination. P-value *** = 0.0001, **** < 0.0001 analyzed via t-test relative to control. (e) DMCs treated with either 25 µM L-DOPA, 75 µM phenylthiourea (PTU), or a combination. P-value **** = 0.0006 analyzed via t-test relative to control. (f) Panel of melanoma cell lines treated with combination 25 µM L-DOPA and 6.25 µM carbidopa. (g) YUMM1.7 murine melanoma growth in syngeneic BL/6 mice treated with vehicle or 300 mg/kg L-DOPA methyl ester and 75 mg/kg carbidopa. P-value ** = 0.0065. n=5 for each group.

To test whether DOPA inhibits MC proliferation, we exposed LMC and DMCs to increasing concentrations of DOPA. DOPA decreased proliferation of LMCs in a dose-responsive and saturable manner, strongly suggesting a specific receptor mediated activity. In contrast, DOPA had no effect on proliferation of DMCs. DOPA effects in LMCs saturated at 6.25 μM. At this exposure, LMCs proliferated at the same rate as DMCs, suggesting that DMCs contain a saturating amount of endogenously synthesized DOPA **(Figure 2c)**. Consistent with this, exogenous DOPA supplementation increased melanin synthesis in LMCs, but did not affect melanin content in DMCs **(Supplemental Figure 2a)**.

To test whether the anti-proliferative effect of DOPA is dependent on melanin synthesis, we utilized the tyrosinase inhibitor N-phenylthiourea (PTU).^32–34^ As tyrosinase catalyzes not only the reaction of tyrosine to DOPA, but also the subsequent conversion of DOPA to dopaquinone **(Figure 2a)**, PTU prevents conversion of exogenous DOPA to melanin. In LMCs, PTU alone had no significant effect on proliferation, while the combination of PTU and DOPA continued to inhibit cell growth **(Figure 2d, Supplemental Figure 2b)**. In DMCs, PTU decreased pigment production and increased proliferation rate. However, DMCs treated with both PTU and DOPA proliferated at the slow baseline DMC rate, even though they remained lightly pigmented **(Figure 2e)**. Together, these data show that DOPA’s effects on MC proliferation are independent of melanin, and that differences in endogenously produced DOPA are likely responsible for most, if not all, of the observed proliferation differences between DMCs and LMCs.

In addition to melanin, the biologic impact of DOPA is also generally attributed to its conversion to dopamine and 3-O-methyldopa (**Figure 2a**), both of which affect activity of dopamine receptors. However, neither of these DOPA metabolites appeared necessary for the anti-proliferative effects of DOPA in MCs. We used the DOPA decarboxylase (DDC) inhibitor, carbidopa, to inhibit the conversion of DOPA to dopamine, and the catechol-O-methyltransferase (COMT) inhibitor, entacapone, to block synthesis of 3-O-methyldopa. Neither inhibitor altered the anti-proliferative effect of DOPA **(Supplemental Figure 2c-e)**. Also consistent with the idea that dopamine is not a mediator of the observed DOPA effects, LC/mass spectrometry analysis did not detect any dopamine in MCs (although we readily detected DOPA in the same samples). To analyze if DOPA impacted other primary cells found in skin, we treated primary human fibroblasts, keratinocytes, and melanocytes with increasing concentrations of DOPA. We observed no effect on primary human keratinocytes, and mild inhibition of fibroblast growth **(Supplemental Figure 2f)**. To test whether melanoma cells also respond to DOPA, we treated multiple human and mouse melanoma cell lines with DOPA/carbidopa and observed marked inhibition of proliferation in most, but not all melanoma lines, independent of BRAF and NRAS mutational status **(Figure 2f, Supplementary Table 1)**. The mechanism responsible the observed DOPA resistance in some lines is determined below, and these lines thereby proved to be quite useful for validating our overall conclusion that DOPA effects are mediated by CHRM1.

We next questioned whether DOPA may have therapeutic utility as a systemically delivered agent for melanoma *in vivo*. Systemic delivery of combined L-DOPA and carbidopa is already FDA-approved for Parkinson’s disease^35^. The DOPA/carbidopa combination is utilized, rather than DOPA alone, because carbidopa inhibits DDC and thereby prevents DOPA from being converted to dopamine everywhere except the brain, where it is needed to treat Parkinson’s: carbidopa does not cross the blood brain barrier, whereas DOPA does. This combination is therefore ideal for our purposes because we wanted to expose the subcutaneous melanomas to DOPA, not to dopamine. BL/6 mice harboring syngeneic YUMM1.7 melanoma (*Braf*^*V600E/wt*^*Pten*^*- /-*^ *Cdkn2*^*-/-*^) were treated with a combination of 300 mg/kg L-DOPA methyl ester and 75 mg/kg carbidopa. Treatment was initiated after tumors reached 3-4 mm in diameter **(Supplemental Figure 2g)**. DOPA/carbidopa treatment was well tolerated by mice and significantly inhibited YUMM1.7 tumor growth **(Figure 2g)**. To understand if the L-DOPA and carbidopa treatment effect in this syngeneic model depends on an immune system response to tumor cells, we repeated the experiment using SCID mice and observed inhibition of tumor growth (**Supplemental Figure 2h**). Together, these results suggest that endogenously synthesized DOPA is the primary determinant of proliferative differences in melanocytes and that exogenous DOPA supplementation inhibits melanoma *in vivo*, independent of an adaptive immune response.

### DOPA antagonizes CHRM1

Data in Figures 1 and 2 strongly suggest that DOPA effects in MCs and melanoma cells are specific and receptor mediated. To identify the receptor(s) responsible, we first considered GPCRs, as melanin synthesis in MCs is classically regulated by the melanocortin receptor MC1R, a Gs-coupled GPCR. To our knowledge, the only previous report associating DOPA with a specific receptor in any cell type identified ocular albinism type 1 (OA1) as a possible DOPA receptor in retina pigment epithelial cells.^36^ To test whether OA1 mediated DOPA effects in melanoma, we depleted OA1 in DOPA sensitive human melanoma cells using siRNA. This had no effect on the DOPA response **(Supplemental Figure 3a)**. To identify new possible GPCR candidates, we used PRESTO-TANGO screening, which is an unbiased high throughput assay to test whether DOPA directly binds to any of the approximately 320 nonolfactory human GPCRs.^37^ We compared top hits to genes expressed in melanocytes and melanoma cells and identified 8 GPCRs predicted to be activated by DOPA, and 9 GPCRs predicted to be inhibited by DOPA **(Figure 3a)**. Simultaneously, we conducted an *in vivo* genetic screen in a human melanoma model using doxycycline-inducible CRISPR-Cas9 to target all non-olfactory human GPCRs **(Figure 3b, Supplemental Figure 4e-g)**. Top hits that appeared in both screens were validated via pooled siRNA knockdown of each GPCR receptor in human A375 melanoma cells. The only siRNA pool that rendered cells insensitive to DOPA was the pool targeting the cholinergic receptor muscarinic 1 (CHRM1), a Gq coupled GPCR **(Figure 3c,d, Supplemental Figure 3a,b)**. These complementary pharmacologic and genetic screens therefore converged upon CHRM1, a GPCR not previously known to bind DOPA, nor to affect melanoma. To further verify results seen with siRNA, we utilized a complementary CRISPR-Cas9 gene editing approach with guide RNAs targeting CHRM1 in A375 melanoma cells. While we were unable to achieve full knockout of CHRM1, potentially due to the hypotriploid karyotype of this model, CRISPR-Cas9 CHRM1 knockdown nonetheless markedly inhibited the antiproliferative effects of DOPA/carbidopa **(Figure 3e, Supplemental Figure 3c)**.

**Figure 3:**
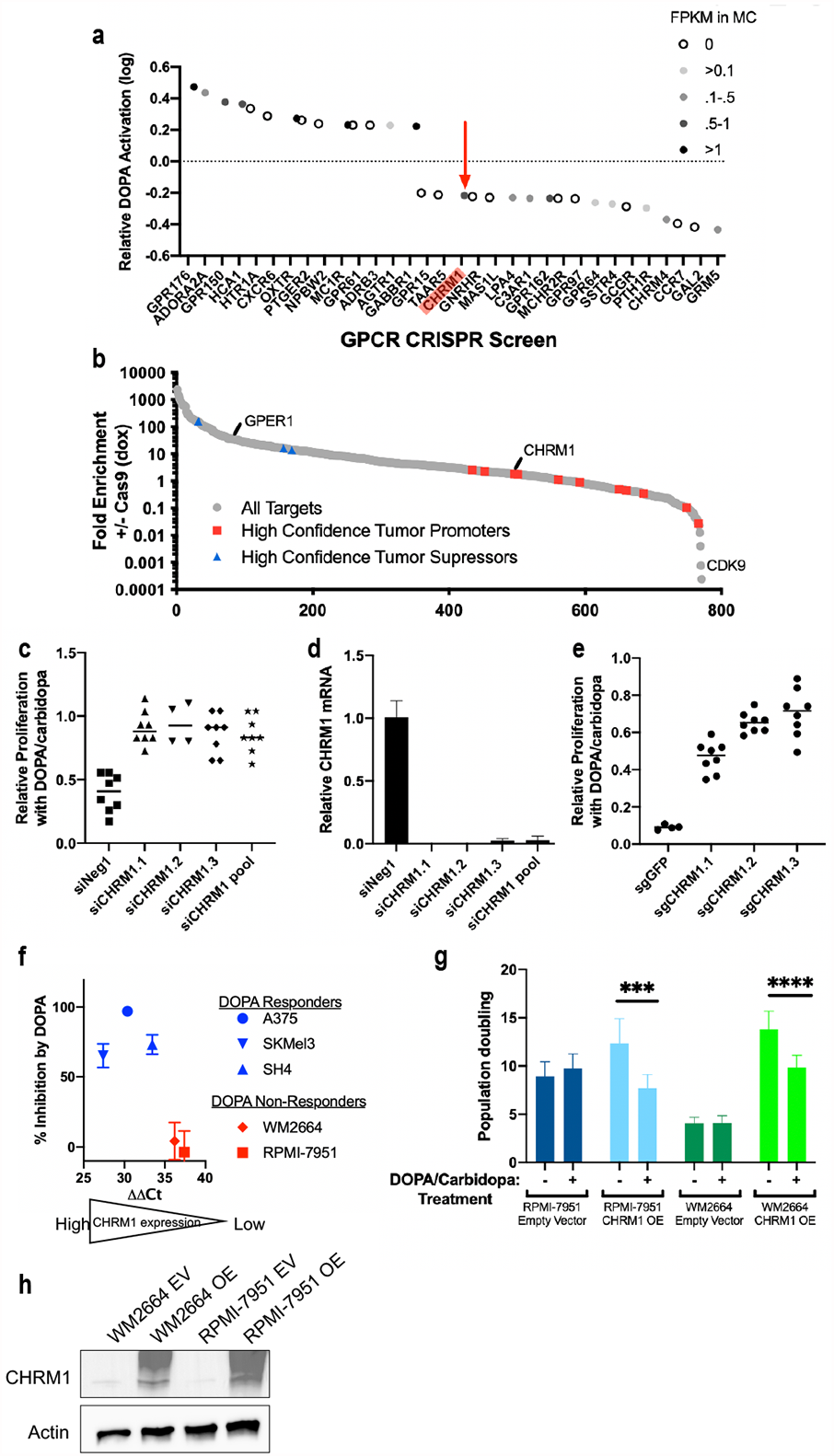
DOPA antagonizes CHRM1. (a) DOPA mediated GPCR activation or inhibition as determined by the PRESTO-Tango reporter assay. Data points are shaded based on relative expression determined using RNA-sequencing in melanocytes (FPKM). (b) Log fold enrichment of CRISPR gRNAs selected for or against. Controls for pro-tumorigenic proteins included CDK9 and PCNA. GPER served as an internal GPCR tumor suppressor control. High confidence hits are targets with at least 5 guides that are selected for (>5-fold) or against (<0.1-fold), and where those 5 guides represent at least 50% of total guides for that gene. (c) siRNA mediated CHRM1depeletion in A375 human melanoma in the presence of 25 µM L-DOPA and 6.25 µM carbidopa after 5 days treatment. (d) qPCR for CHRM1 mRNA in A375 after siRNA treatment confirming knockdown. Timepoint taken 24 hours after siRNA transfection. (e) Effect of 25 µM L-DOPA and 6.25 µM carbidopa on proliferation of A375 cells in which CHRM1 was depleted using CRISPR-Cas9 vs control gRNA against GFP. Cell number was determined at day 5. (f) Low CHRM1 expression, determined via qPCR, correlates with lack of response to 25 µM L-DOPA and 6.25 µM carbidopa. (g) CHRM1 overexpression in WM2664 and RPMI-7951 human melanoma (DOPA non-responders) in the presence or absence of 25 µM L-DOPA and 6.25 µM carbidopa after 5 days treatment. P-value *** = 0.0002, ****<0.0001 analyzed via t-test. (h) Western blot for CHRM1 in WM2664 and RPMI-7951 after transduction with either empty vector or CHRM1.

Consistent with these data showing that CHRM1 mediates DOPA effects, DOPA responses in melanoma cell lines correlated with CHRM1 expression, with DOPA insensitive cells lacking CHRM1 expression **(Figure 3f)**. As DOPA appeared to function as a CHRM1 antagonist, we next tested whether the known CHRM1 synthetic antagonist, pirenzepine^38,39^, mimics the observed DOPA effects. In a dose dependent manner, pirenzepine recapitulated the anti-proliferative effects of DOPA/carbidopa treatment in A375 human melanoma, but not WM2664, as these cells do not express CHRM1. In contrast, the CHRM1 agonist, pilocarpine^40,41^, had opposite effects, and promoted proliferation in both melanoma cells and DMCs **(Supplemental Figure 3d-f)**. The endogenous agonist of CHRM1 is acetylcholine (ACh); although we did not detect ACh in primary MC cultures *in vitro*, ACh from nonneuronal sources is abundant in human skin.^42–45^ ACh promoted proliferation of DMCs, but not LMCs, and this effect was inhibited by DOPA treatment (**Supplemental Figure 3g,h**). Together these data show that CHRM1 activation promotes MC and melanoma cell proliferation, and that CHRM1 is necessary for the anti-proliferative effects of DOPA.

To determine if CHRM1 expression is sufficient to confer DOPA sensitivity to DOPA insensitive melanoma cells lacking CHRM1, we used lentiviral transduction to express CHRM1 in two non-responding melanoma cell lines, RPMI-7951 and WM2664. Upon CHRM1 expression, cells grew faster than parental controls, suggesting that CHRM1 may function as a melanoma oncodriver **(Figure 3g**,**h)**. Consistent with this idea, analysis of 9,736 tumors and 8,587 normal samples from the TCGA and the GTEx projects^46^, shows that high CHRM1 expression in melanoma is associated with decreased overall survival and increased stage progression **(Supplemental Figure 4a, b)**. Importantly, CHRM1 expression rendered RPMI-7951 and WM2664 newly sensitive to DOPA, supporting the idea that CHRM1 is both necessary and sufficient for DOPA effects in MC and melanoma **(Figure 3g**,**h)**. To further confirm the specificity of these genetic and pharmacologic data, and to control for possible off target effects of the CHRM1 targeting gRNA, we used lentiviral transduction to restore CHRM1 expression in A375 cells in which we had previously depleted CHRM1 using CRISPR-Cas9. With this transgene rescue, cells were resensitized to DOPA **(Supplemental Figure 4c**,**d)**. Together, these data show that CHRM1 is the major mediator of DOPA effects in melanoma.

### DOPA inhibits oncogenic Gq signaling and represses FOXM1

CHRM1 is a Gq coupled GPCR known to activate both Ras/MAPK and PI3K/Akt signaling in other cell types.^47–49^ Both of these pathways are major drivers of melanoma and other cancers, and are targets of approved inhibitors used clinically.^50–53^ Consistent with our discovery that CHRM1 is a DOPA sensitive melanoma driver, exogenous DOPA induced rapid depletion of both phosphorylated extracellular-signal-regulated kinase (ERK), and phosphorylated AKT in melanoma cells **(Supplemental Figure 5a)**. We also noted parallel depletion of c-Myc and FOXM1, which both function as transcription factors and proliferation drivers positively regulated by MAPK and AKT ^54–58^ **(Figure 4a, Supplemental Figure 5b)**. Importantly, we also found that LMCs, which synthesize less endogenous DOPA than DMCs, contain higher levels of FOXM1 protein **(Figure 4b)**.

**Figure 4:**
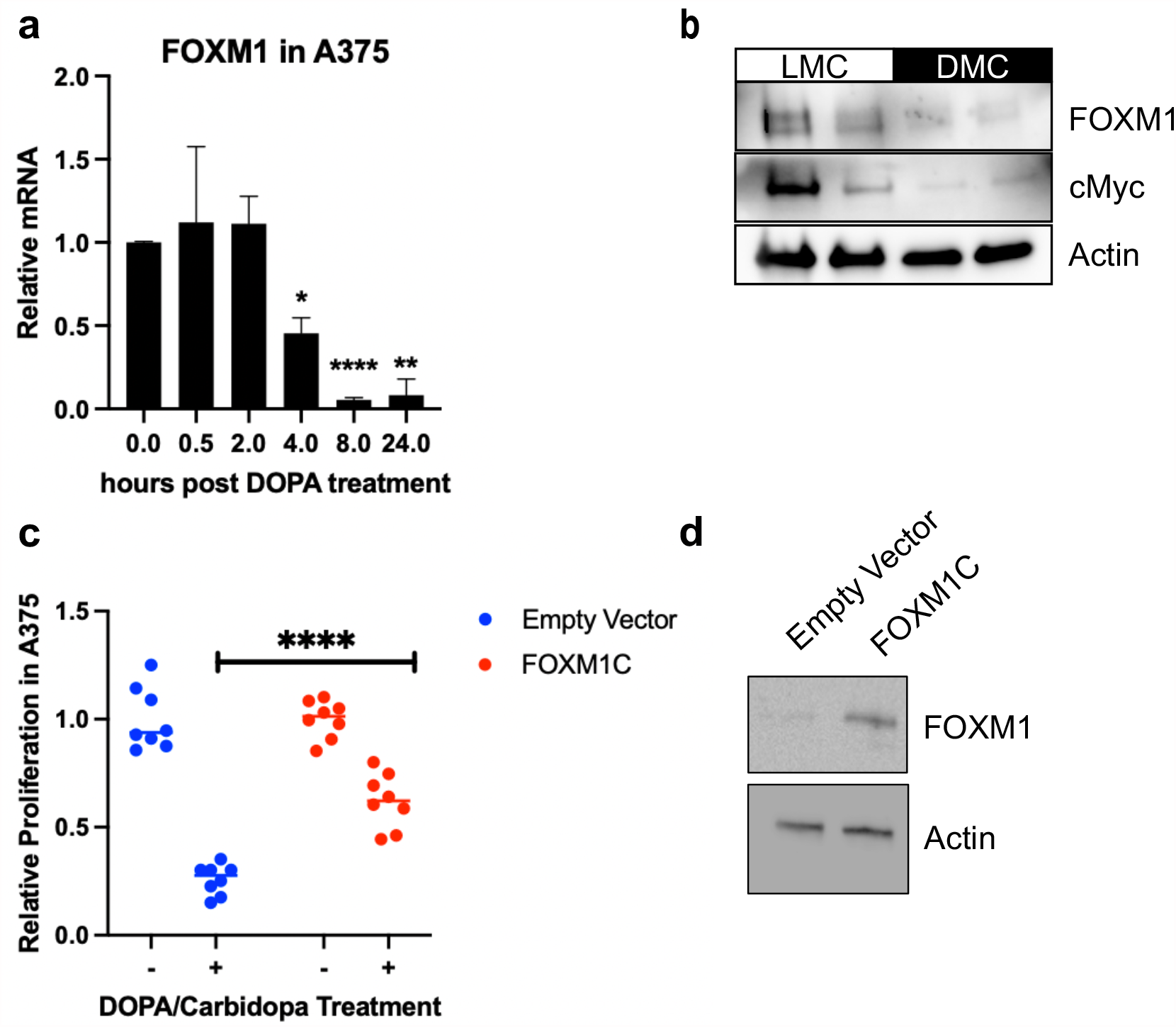
DOPA inhibits oncogenic Gq signaling and represses FOXM1. (a) FOXM1 mRNA-level determined via qPCR of time-course in A375 human melanoma treated with 25 µM L-DOPA and 6.25 µM carbidopa for increasing amounts of time. P-value * = 0.0142, ** = 0.0054, **** < 0.0001. (b) Western blot of FOXM1 and c-Myc at baseline in light and dark melanocytes. (c) Proliferation in A375 cells following transduction with FOXM1C versus empty vector +/- 25 µM L-DOPA and 6.25 µM carbidopa. P-value **** < 0.0001. (d) Western blot confirming FOXM1C overexpression in A375 human melanoma.

We were specifically interested in this DOPA-induced FOXM1 depletion, as FOXM1 is overexpressed in up to 70% of metastatic melanomas and high expression correlates with worse outcomes.^55,59,60^ FOXM1 stimulates cell growth by promoting genes critical for cell proliferation and is a key regulator of the G1-S phase transition. To test whether FOXM1 loss was necessary for the anti-proliferative effects of DOPA, we overexpressed FOXM1C, the primary isoform in melanocytes and melanoma.^55^ This attenuated, but did not completely abolish, DOPA’s anti-proliferative effect **(Figure 4c**,**d)**.

### Pharmacologic inhibition of FOXM1 suppresses melanoma growth

Encouraged by our data showing that FOXM1 loss downstream of CHRM1 was necessary for the anti-proliferative effects of DOPA, we next questioned whether FOXM1 inhibition alone was sufficient to similarly inhibit melanoma proliferation **(Figure 5a)**. Historically, transcription factors have been viewed as generally undruggable targets.^61^ However, small molecule inhibitors have recently been developed for FOXM1 that block DNA binding.^62^ In vitro exposure to the FOXM1 inhibitor FDI-6 markedly reduced melanoma cell proliferation and, most strikingly, included a dramatic change in melanoma cell morphology: cells became multipolar and larger, and generally appeared more like normal primary melanocytes than the untreated melanoma cells, which had a rounded/oval appearance **(Figure 5b**,**c, Supplemental Figure 5c)**. These morphologic features have also been recognized by others as indicative of a more fully differentiated melanocyte cell state.^63,64^ Consistent with this idea, LMCs were more sensitive to FDI-6 treatment than DMCs **(Supplemental Figure 5d)**.

**Figure 5:**
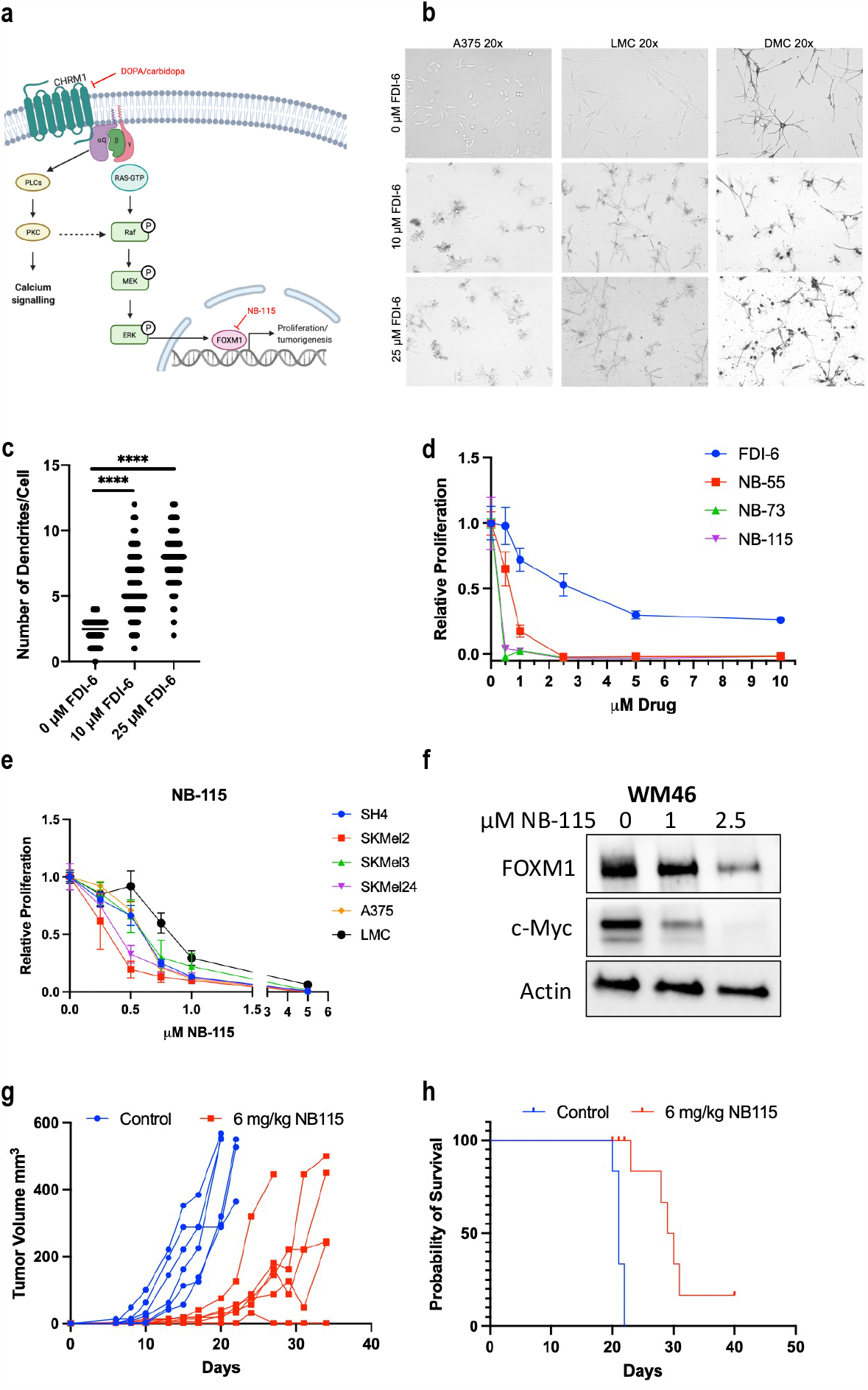
Pharmacologic FOXM1 inhibition suppresses melanoma growth and extends animal survival. (a) Schematic overview of CHRM1 signaling in melanoma highlighting drugs used in this paper to inhibit the pathway. (b) Morphologic appearance of A375 human melanoma, lightly pigmented melanocytes (LMC), and darkly pigmented melanocytes (DMC) after 24 hour exposure to increasing concentrations of FDI-6 (FOXM1i). P-value **** < 0.0001 analyzed via t-test. (c) Change in number of dendrites per A375 cell after exposure to FDI-6 for 24 hours. 10 representative fields at 10x magnification from each condition were quantified. (d) Proliferation of A375 human melanoma cells in presence of increasing concentrations of FOXM1 inhibitors, including FDI-6 (commercially available), NB-55, NB-73, and NB-115. Cell proliferation assay using WST-8 cell viability dye. n=5. (e) Proliferation of a panel of melanoma cell lines in presence of increasing concentrations of NB-115.. n=5. (f) FOXM1 and c-Myc protein in WM46 human melanoma after exposure to NB-115 for 24 hours. (g) YUMM1.7 melanoma growth over time in BL/6 mice treated with vehicle or 6 mg/kg NB-115. N=6 for each group across two identical experiments. (h) Survival probability over time of mice treated with vehicle or 6 mg/kg NB-115. N=6 for each group across two identical experiments. p-value **** < 0.0001 by Mantel-Cox test.

While FDI-6 shows promising results *in vitro* and is a useful and readily available research tool, it has very poor pharmacokinetic properties and is not useful for in vivo studies.^65^ However, a new class of FOXM1 inhibitors was recently shown to have activity in preclinical breast cancer models, without significant systemic toxicity.^65^ Three of these new FOXM1 inhibitors, NB-55, NB-73, and NB-115, were more effective than FDI-6 at inhibiting melanoma proliferation **(Figure 5d)**. Consistent with the idea that FOXM1 is a critical element downstream of CHRM1, NB-115 inhibited cell growth in a variety of human and mouse melanoma cell lines, including those that do not respond to DOPA because they lack CHRM1. NB-115 mediated FOXM1 depletion was associated with depletion of FOXM1 protein itself, as well as depletion of c-Myc **(Figure 5e**,**f, Supplemental Figure 5e**,**f)**. FOXM1 and c-Myc are both known to positively regulate the transcription of each the other ^66–68^, and the observed loss of FOXM1 agrees with previous reports establishing that NB-55, NB-73, and NB-115 promote proteasome mediated FOXM1 degradation.^65^

We next tested if systemically delivered NB-115 inhibited melanoma in vivo. BL/6 mice harboring syngeneic YUMM1.7 melanoma (*Braf*^*V600E/wt*^*Pten*^*-/-*^ *Cdkn2*^*-/-*^) were treated with 6 mg/kg NB-115. This significantly inhibited YUMM1.7 melanoma growth and extended overall survival, with one mouse completely clearing its tumor **(Figure 5g**,**h)**. Together, these data suggest that CHRM1 is a melanoma target that is regulated via DOPA, which is naturally synthesized in melanocytes. Further, FOXM1 is a critical downstream regulator of DOPA’s anti-proliferative effect and itself appears to be a potential therapeutic target.

## Discussion

Decades of clinical and epidemiological data suggest that the physical UV shielding effect of is insufficient to fully explain the difference in melanoma incidence between lightly and darkly pigmented skin. We hypothesized that the mechanisms responsible for this also contribute to the differences in proliferation rate that we routinely observe between LMC and DMCs. To our knowledge, this is the first work to directly explore the mechanism responsible for these differences, to show that CHRM1 is a DOPA receptor, to show that CHRM1 affects melanocyte homeostasis, and to demonstrate that CHRM1 and FOXM1 are potential therapeutic targets for melanoma. Future research utilizing melanocytes, as well as melanoma clinical trials, may benefit from consideration of the genetic background of the cells, as differences in DOPA and CHRM1 signaling are likely to affect some of the experimental results and interpretation.

Our data are consistent with some provocative, but mechanistically unexplained findings from older literature. DOPA was shown 28 years ago to bind to a protein in rodent melanoma cell membranes although the specific protein was not identified, and the functional consequences of that binding for MC function or melanoma pathology were not determined.^69,70^ Additionally, studies have identified L-DOPA as a regulator of melanocyte functions, although the mechanism(s) responsible were not established.^71,72^ Even more tantalizing, 43 years ago, L-DOPA methyl ester was shown to inhibit B16 melanoma in mice, but whether that resulted from DOPA itself, melanin, or other metabolite, was not determined. Most critically, the receptor and signaling mechanism(s) mediating that DOPA effect were not determined, and this old data appears to be mostly forgotten in recent melanoma literature.^73–78^

Acetylcholine, which is abundantly available in human skin^42,43^, signals through the muscarinic acetylcholine receptors (mAChRs), including CHRM1, and these receptors have been shown to be present in normal human melanocytes.^79^ Signaling through mAChRs impacts a wide spectrum of diseases and, as such, many mAChRs antagonists are already approved in the U.S for use in people. Among these are atropine for childhood myopia^80^ and scopolamine for motion sickness^81^. Unfortunately, these agents have very short half lives in vivo, and are thereby not suitable for cancer studies. Nonetheless, we have shown that future mAChR antagonists with improved systemic pharmacokinetic properties may be effective against melanoma. Although, cholinergic muscarinic receptors are best known for their activity in the nervous system, ours is not the first work to implicate ACh in cancer progression, as recent work in murine prostate cancer models established that the nerves activate pro-tumorigenic cholinergic signaling in the tumor microenvironment that promotes tumor invasion and metastasis.^82^

We showed that combination treatment of DOPA and carbidopa, an FDA approved therapy for Parkinson’s disease, mimics the effects of endogenously produced DOPA in DMCs, and thereby inhibits melanoma. Parkinson’s disease is a neurodegenerative disorder caused by a loss of neurons in the substantia nigra, ultimately leading to dopamine deficiency that quickly leads to a decline in motor function.^83^ Multiple epidemiological studies have found an association between melanoma and Parkinson’s disease melanoma.^84–86^ This association is reciprocal: patients with melanoma have an increased risk of Parkinson’s disease and patients with Parkinson’s disease are more likely to develop melanoma. Studies have also shown that incidence in Parkinson’s disease is 2-3 times more common in white populations as compared to African-American populations,.^87–89^ These epidemiological studies, together with this current work, suggest that the relative lack of DOPA in lightly pigmented individuals may predispose them not only to melanoma, but also Parkinson’s; however, the pathobiology of Parkinson’s disease is complex and further investigation is needed to determine whether these two seemingly disparate diseases are mechanistically linked through DOPA.

Finally, we established that pharmacologic DOPA/carbidopa led to decreased activation of both the MAPK and AKT pathways and ultimately downregulation of FOXM1, a major cancer driver.^60,90^ While FOXM1 is downstream of both the MAPK and AKT pathways, FOXM1 depletion is unlikely to be the sole mechanism by which DOPA inhibits melanoma as FOXM1 overexpression only partially rescued cell proliferation in the face of exogenous DOPA. However, selective pharmacologic FOXM1 inhibition significantly inhibited melanoma cell growth *in vitro* and *in vivo*, suggesting it may also be a new melanoma therapeutic target. Future studies will be needed to determine whether the utility of this new class of FOXM1 inhibitors extends to non-cutaneous melanoma and other cancers.

Together, this work demonstrates how the natural genetic diversity in people can be used as a window to discover new signaling pathways regulating normal tissue homeostasis and carcinogenesis.

## Materials and Methods

### Cell culture and proliferation assays

Primary human melanocytes, keratinocytes, and fibroblasts were extracted from fresh discarded human foreskin and surgical specimens as previously described.^28,91–93^ Keratinocytes were cultured in a 1:1 mixture of Gibco Keratinocytes-SFM medium + L-glutamine + EGF + BPE and Gibco Cascade Biologics 154 medium with 1% penicillin-streptomycin (Thermo Fisher Scientific. # 15140122). Fibroblasts were cultured in DMEM (Mediatech, Manassas, VA, USA) with 5% FBS (Invitrogen, Carlsbad, CA, USA) and 1% Penicillin-Streptomycin. Primary melanocytes and human-engineered melanoma cells (heMel) were cultured in Medium 254 (ThermoFisher, #M254500) with 1% penicillin-streptomycin.

YUMM1.7, SH-4 and SK-MEL-2 cells were purchased from ATCC (YUMM1.7 ATCC® CRL-3362™; SH-4 ATCC® CRL-7724™; SK-MEL-2 ATCC® HTB-68™) and cultured in DMEM with 5% FBS and 1% Penicillin-Streptomycin. SK-MEL-3 cells were purchased from ATCC (ATCC® HTB-69™ and cultured in McCoy’s 5A (Modified) Medium with 15% FBS (Invitrogen, Carlsbad, CA, USA) and 1% Penicillin-Streptomycin. RPMI-7951 and SK-MEL-24 cells were purchased from ATCC (RPMI-7951 ATCC® HTB-66™; ATCC® HTB-71™) and cultured in Eagle’s Minimum Essential Medium with 15% FBS and 1% Penicillin-Streptomycin. WM46 and WM2664 melanoma cells were a gift from Meenhard Herlyn (Wistar Institute, Philadelphia, PA, USA) and were cultured in TU2% media. Tumor cells were regularly tested using MycoAlert Mycoplasma Detection Kit from Lonza (Allendale, NJ, USA).

For monitoring cell proliferation 10×10^5^ YUMM1.7 or A375, 12×10^5^ RPMI-7951, 15×10^5^ WM46, WM2664, SH4, SK-MEL-2, SK-MEL-24, SK-MEL-3, or 30×10^5^ melanocytes were seeded per well in 12-well cell culture plates. Cells were treated every second day and manually counted in triplicate using a hemocytometer. All the experiments were performed in cell populations that were in culture during a maximum of 3 weeks (5 passages in average) since thaw from the corresponding stock.

3,4-Dihydoxy-L-phenylalanine (D9628), N-Phenylthiourea (P7629), and FDI-6 (SML1392) were purchased from Sigma-Aldrich (St. Louis, MO, USA). (S)-(-)-Carbidopa (0455), Pirenzepine dihydrocholoride (1071), Pilocarpine hydrocholride (0694) were purchased from Tocris Bioscience (Bristol, United Kingdom). 3-O-methyl-L-DOPA hydrate (20737) and Entacapone (14153) were purchased from Cayman Chemicals (Ann Arbor, MI, USA). NB-55, NB-73, and NB-115 were prepared as described.^65^

### Genetic manipulation of CHRM1

We used lentiviral transduction to deliver dox-inducible Cas9 and gRNA targeting CHRM1 in human A375 melanoma cells. Three different guide RNAs were used to target CHRM1. Transduced cells were selected with puromycin, and single cells subsequently isolated, expanded and examined for CHRM1 protein expression, compared to clones isolated in parallel with no doxycycline treatment. The following gRNA sequences were used (5’-3’):

sgCHRM1.1_Fw: caccgGCTCCGAGACGCCAGGCAAA

sgCHRM1.1_Rv: aaacTTTGCCTGGCGTCTCGGAGCc

sgCHRM1.2_Fw: caccgGATGCCAATGGTGGACCCCG

sgCHRM1.2_Rv: aaacCGGGGTCCACCATTGGCATCc

sgCHRM1.3_Fw: caccgCAAGCGGAAGACCTTCTCGC

sgCHRM1.3_Rv: aaacGCGAGAAGGTCTTCCGCTTGc

Using ThermoFisher’s Silencer Silect protocol, we knocked down CHRM1 in human A375 melanoma cells. Briefly, each siRNA was diluted in Opti-MEM medium (Invitrogen, 31985062) to a concentration of 10 µM, to ultimately be diluted to 30 pmol in a 6-well plate. If siRNA were pooled, each individual siRNA was used at 10 pmol (for a combined total of 30 pmol) in a 6-well plate. Diluted siRNA’s were combined with diluted Lipofectmaine (Invitrogen, 11668027) and incubated on cells for 24 hours. After 24 hours cells were plated in a 12 well plate with 10,000 cells per well and treated with a combination of DOPA and carbidopa for four days.

We used three different siRNA against CHRM1: s3023 (labeled siCHRM1.1), s3024 (labeled siCHRM1.2), s553080 (labeled siCHRM1.3). Negative controls included Negative Control No.1 (ThermoFisher, 4390843) and Negative Control No. 2 (ThermoFisher, 4390846) and a positive control against Kif11 (University of Pennsylvania, High-throughput sequencing core).

### Human-engineered melanoma xenografts

Organotypic skin grafts were established using modifications to previously detailed methods.^28,94^ The Keratinocyte Growth Media (KGM) used for keratinocyte-only skin grafts was replaced with modified Melanocyte Xenograft Seeding Media (MXSM). MXSM is a 1:1 mixture of KGM, lacking cholera toxin, and Keratinocyte Media 50/50 (Gibco) containing 2% FBS, 1.2 mM calcium chloride, 100 nM Et-3 (endothelin 3), 10 ng/mL rhSCF (recombinant human stem cell factor), and 4.5 ng/mL r-basic FGF (recombinant basic fibroblast growth factor). Briefly, primary human melanocytes were transduced with lentivirus carrying BRAF(V600E), dominant-negative p53(R248W), active CDK4(R24C) and hTERT. Transduced melanocytes (1 × 10^5^ cells) and keratinocytes (5 × 10^5^ cells) were suspended in 80 μL MXSM, seeded onto the dermis, and incubated at 37°C for 4 days at the air–liquid interface to establish organotypic skin. Organotypic skin tissues were grafted onto 5–7-week-old female ICR SCID mice (Taconic) according to an IACUC–approved protocol at the University of Pennsylvania. Mice were anesthetized in an isoflurane chamber and murine skin was removed from the upper dorsal region of the mouse. Organotypic human skin was reduced to a uniform 11 mm × 11 mm square and grafted onto the back of the mouse with individual interrupted 6–0 nylon sutures. Mice were dressed with Bactroban ointment, Adaptic, Telfa pad, and Coban wrap. Dressings were removed 2 weeks after grafting. Mice were sacrificed 100 days after grafting and organotypic skin was removed for histology.

### Subcutaneous tumors and treatments

All mice were purchased from Taconic Biosciences, Inc. (Rensselaer, NY, USA). These studies were performed without inclusion/exclusion criteria or blinding but included randomization. Based on a two-fold anticipated effect, we performed experiments with at least 5 biological replicates. All procedures were performed in accordance with International Animal Care and Use Committee (IACUC)-approved protocols at the University of Pennsylvania. Subcutaneous tumors were initiated by injecting 10 × 10^5^ YUMM1.7 cells in 50% Matrigel (Corning, Bedford, MA, USA) into the subcutaneous space on the left and right flanks of mice. For L-DOPA and carbidopa experiments, L-DOPA methyl ester (300 mg/kg, Tocris # 0455) and carbidopa (75 mg/kg, Cayman #16149) were injected intraperitoneally daily for three weeks, then five days on, two days off for the remainder of the experiment. In the SCID mouse experiment, drugs were injected three days on, one day off for the entire experiment. Both drugs were resuspended in normal saline. Adhering to previous literature^95,96^, carbidopa was injected one hour before L-DOPA injection. For FOXM1 inhibitor experiments, NB-115 (6 mg/kg) was injected subcutaneously for five days on, two days off. NB-115 was dissolved in DMSO and diluted 1:10 in sesame oil to form a stable, homogenous suspension. As subcutaneous tumors grew in mice, perpendicular tumor diameters were measured using calipers. Volume was calculated using the formula L × W^2^ × 0.52, where L is the longest dimension and W is the perpendicular dimension. Animals were euthanized when tumors exceeded a protocol-specified size of 500 mm^3^. Secondary endpoints include severe ulceration, death, and any other condition that falls within the IACUC guidelines for Rodent Tumor and Cancer Models at the University of Pennsylvania.

### Western blot analysis

Adherent cells were washed once with DPBS, and lysed with 8M urea containing 50 mM NaCl and 50 mM Tris-HCl, pH 8.3, 10 mM dithiothreitol, 50 mM iodoacetamide. Lysates were quantified (Bradford assay), normalized, reduced, and resolved by SDS gel electrophoresis on 4–15% Tris/Glycine gels (Bio-Rad, Hercules, CA, USA). Resolved protein was transferred to PVDF membranes (Millipore, Billerica, MA, USA) using a Semi-Dry Transfer Cell (Bio-Rad), blocked in 5% BSA in TBS-T and probed with primary antibodies recognizing β-Actin (Cell Signaling Technology, #3700, 1:4000, Danvers, MA, USA), c-Myc (Cell Signaling Technology, #5605, 1:1000), p-RB (Cell Signaling Technology, #8516, 1:1000), RB (Cell Signaling Technology, #9313, 1:1000), p-CREB (Cell Signaling Technology, #9198, 1:1000), CREB (Cell Signaling Technology, #9104, 1:1000), tyrosinase (Abcam, T311, 1:1000), p53 (Cell Signaling Technology, #2527, 1:1000), CDK4 (Cell Signaling Technology, #12790, 1:1000), P-ERK (Cell Signaling Technology, Phospho-p44/42 MAPK (Erk1/2) (Thr202/Tyr204) (D13.14.4E) XP® Rabbit mAb #4370. 1:1000), ERK (Cell Signaling Technology, p44/42 MAPK (Erk1/2) (137F5) Rabbit mAb #4695, 1:1000), pAKT S473 (Cell Signaling Technology, #9271, 1:1000), AKT (Cell Signaling Technology, #9272, 1:1000) CHRM1 (Invitrogen, #PA5-95151, 1:1000), FoxM1 (Cell Signaling Technology, #5436, 1:1000). After incubation with the appropriate secondary antibody, proteins were detected using either Luminata Crescendo Western HRP Substrate (Millipore) or ECL Western Blotting Analysis System (GE Healthcare, Bensalem, PA). After incubation with the appropriate secondary antibody [(Rabbit Anti-Mouse IgG H&L (Biotin) preabsoFS9rbed (ab7074); Anti-mouse IgG, HRP-linked Antibody #7076. 1:2000)] proteins were detected using ClarityTM Western ECL Substrate (Bio-Rad. #170-5060). All western blots were repeated at least 3 times.

### Quantitative RT-PCR

RNA was extracted using RNeasy kit (Qiagen. #74104) following the manufacturer’s instructions. cDNA was obtained using High Capacity cDNA Reverse Transcription Kit (Applied Biosystems #4368814). For quantitative real-time PCR, PowerUP™ SYBR™ Green Master Mix (Applied Biosystems #A25741) was used. ViiA 7 Real-Time PCR System was used to perform the reaction (Applied Biosystems). Values were corrected by b-actin expression. The 2^−ΔΔCt^ method was applied to calculate the relative gene expression. Primers used are included in Table S2.

### PRESTO-TANGO

We used the National Institute of Mental Health’s Psychoactive Drug Screening Program at the University of North Carolina to perform PRESTO-TANGO^37^ analysis of 400 non-olfactory GPCRs in the presence or absence of L-DOPA.

### In vivo CRISPR Screen

We used lentiviral transduction to deliver dox-inducible Cas9 to WM46 cells and pulled tightly controlled clones and verified by western blot. The non-olfactory GPCR CRISPR library was transduced with lentivirus with a MOI less than 1. 1,000,000 cells were injected subcutaneously in SCID mice. After 7 days of tumor formation, mice were fed dox chow to activate Cas9. After 56 days, tumors were harvested and frozen for sequencing.

Genomic DNA was extracted, and 30 independent PCR reactions used to amplify the sgRNA sequences (100ng DNA/reaction). Pooled PCR products were prepared for library construction and sequencing via MiSeq (Illumina).

Demultiplexed FASTQ files were processed using cutadapt 1.15. The number of reads for each sgRNA was estimated using the MAGeCK 0.5.7 count module. Reads for each sgRNA were normalized as follows:

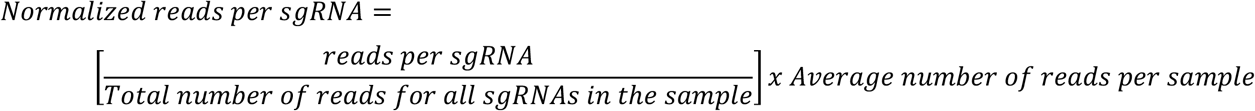

If a given sgRNA was not represented in two or more control tumors (*i*.*e*., tumors that were not subject to dox selection), we removed the sgRNA from our downstream analysis. Normalized reads for each sgRNA were averaged over each condition (+ dox and – dox) and the fold change (FC) was calculated as:

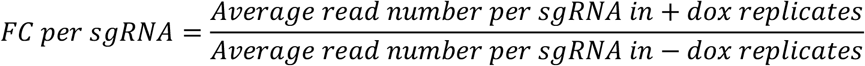

For dropout hit identification we chose genes targeted by more than or equal to 2 sgRNAs that show a fold change of at least 0.1. Genes were ranked based on the average FC of all represented sgRNA targeting the gene.

### Immunohistochemistry and quantification

Formalin-fixed paraffin-embedded (FFPE) human skin tissue sections from organotypic tissue was stained for MITF (NCL-L-MITF, Leica Biosystems, Nussloch, Germany), MelanA (NCL-L-MITF, Leica Biosystems), and Ki67 (NCL-L-Ki67-MM1, Leica Biosystems). Staining was performed following the manufacturer’s protocol for high temperature antigen unmasking technique for paraffin sections. For melanin staining, FFPE embedded tissue was subjected to Fontana-Masson histochemical stain as previously described.^18,19^

Tissue section quantification was performed according to previous reports.^97^ Briefly, 10X photomicrograph images of representative tissue sections were taken using the Keyence BZ-X710 (Itasca, IL, USA). Tiff files of the images were saved and transferred to FIJI (Image J). Images corresponding to the single specific color were then analyzed to determine the number of pixels in each sample and normalized to epidermal area. The numbers of pixels representing Melan-A staining were normalized to the total amount of epidermal area.

### HPLC-MS

We used the Metabolomics Core at the Children’s Hospital of Philadelphia (https://www.research.chop.edu/metabolomic-core) to perform HPLC for DOPA in lightly and darkly pigmented melanocytes. Cells were scraped from tissue culture plates, resuspended in 4% perchloric acid (PCA), and immediately brought to core.

### Melanin assay

Cells (1 × 10^5^) were seeded uniformly on 6-well tissue culture plates. Cells were treated with vehicle controls, DOPA, or PTU for 7 days. Cells were then trypsinized, counted, and pellets containing 300,000 cells were spun at 300 g for 5 min. The cell pellets were solubilized in 120 μL of 1M NaOH, and boiled at 100C for 5 min. The optical density of the resulting solution was read at 450 nm using an Emax microplate reader (Molecular Devices, Sunnyvale, CA, USA). The absorbance was normalized to a control pellet of 300,000 WM46 cells. All melanin assays were repeated at least three times and each time performed in triplicate.

### Statistical analysis

All statistical analysis was performed using Graphpad Prism 8 (Graphpad Software, La Jolla, CA, USA). No statistical methods were used to predetermine sample size. Details of each statistical test used are included in the figure legends.

## Acknowledgements

The authors thank the University of Pennsylvania Skin Biology and Disease Research -based center for analysis of tissue sections. GPCR functional data was generously provided by the National Institute of Mental Health’s Psychoactive Drug Screening Program, Contract # HHSN-271-2018-00023-C (NIMH PDSP). The NIMH PDSP is Directed by Bryan L. Roth MD, PhD at the University of North Carolina at Chapel Hill and Project Officer Jamie Driscoll at NIMH, Bethesda MD, USA.

## Author Contributions

**M.D**. designed and conducted experiments; performed data collection and analysis; conceptualized and supervised the project; wrote and reviewed the manuscript. **C.A.N**. designed and conducted experiments; conceptualized and reviewed the manuscript. **I.Y.L**. conducted experiments; reviewed the manuscript. **S.V**. performed data analysis. **S.H.K**. provided methodology and synthesized resources. **J.A.B**. provided methodology and resources; reviewed the manuscript. **B.S.K**. provided methodology and resources; reviewed the manuscript. **M.E.F**. conceptualized experiments; provided resources; performed data analysis; reviewed the manuscript. **T.W.R**.: conceptualized and supervised the project; reviewed the manuscript.

---

**Supplemental Figure 1:**
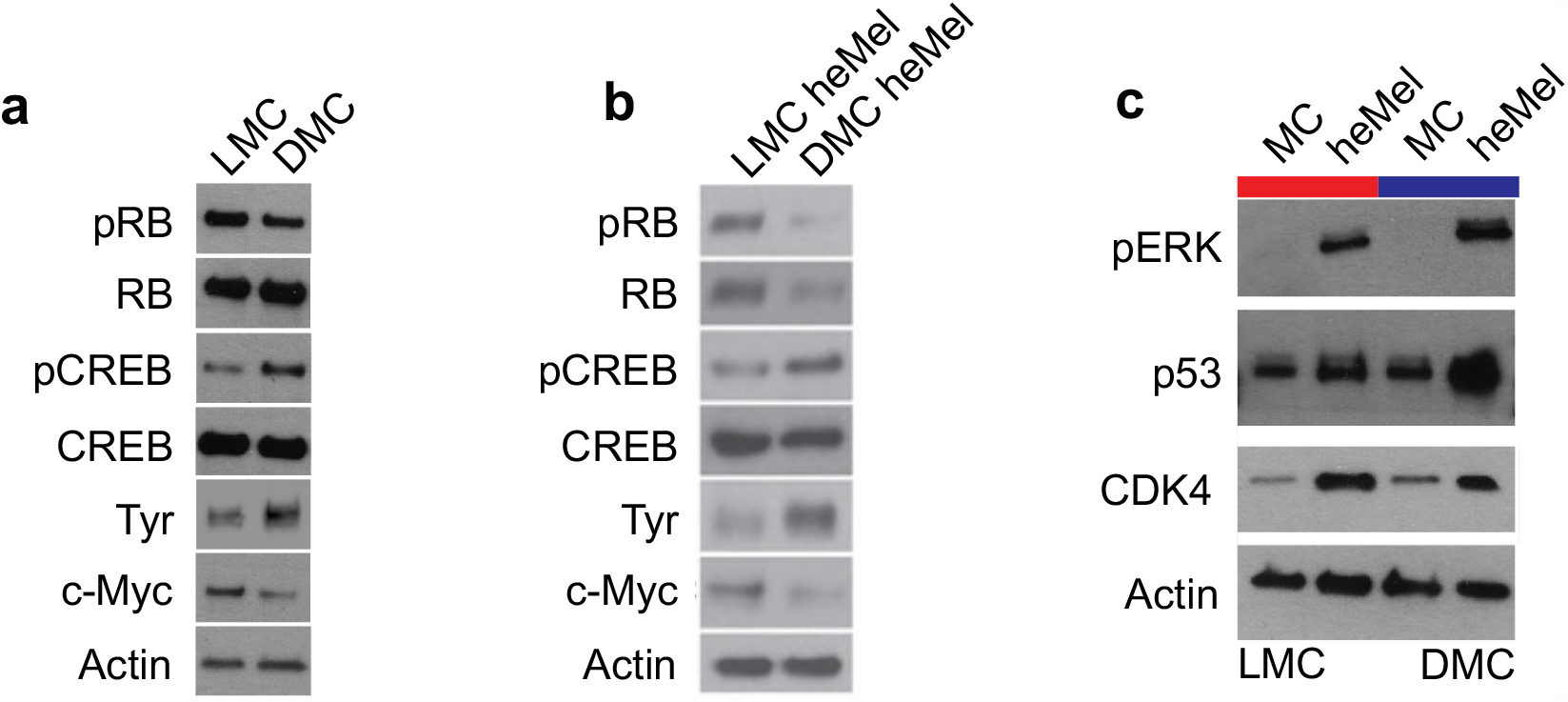
LMCs express lower levels of classical melanocyte differentiation markers, and higher levels of the proliferation driver and stem cell marker c-Myc than DMCs. (a) Western blot of differentiation markers in representative lightly pigmented melanocytes (LMC) and darkly pigmented melanocytes (DMC). (b) Western blot of differentiation markers in representative lightly pigmented heMel (LMC heMel) and darkly pigmented heMel (DMC heMel). (c) Western blot of LMC and DMC transduced with heMel overexpression vectors.

**Supplemental Figure 2:**
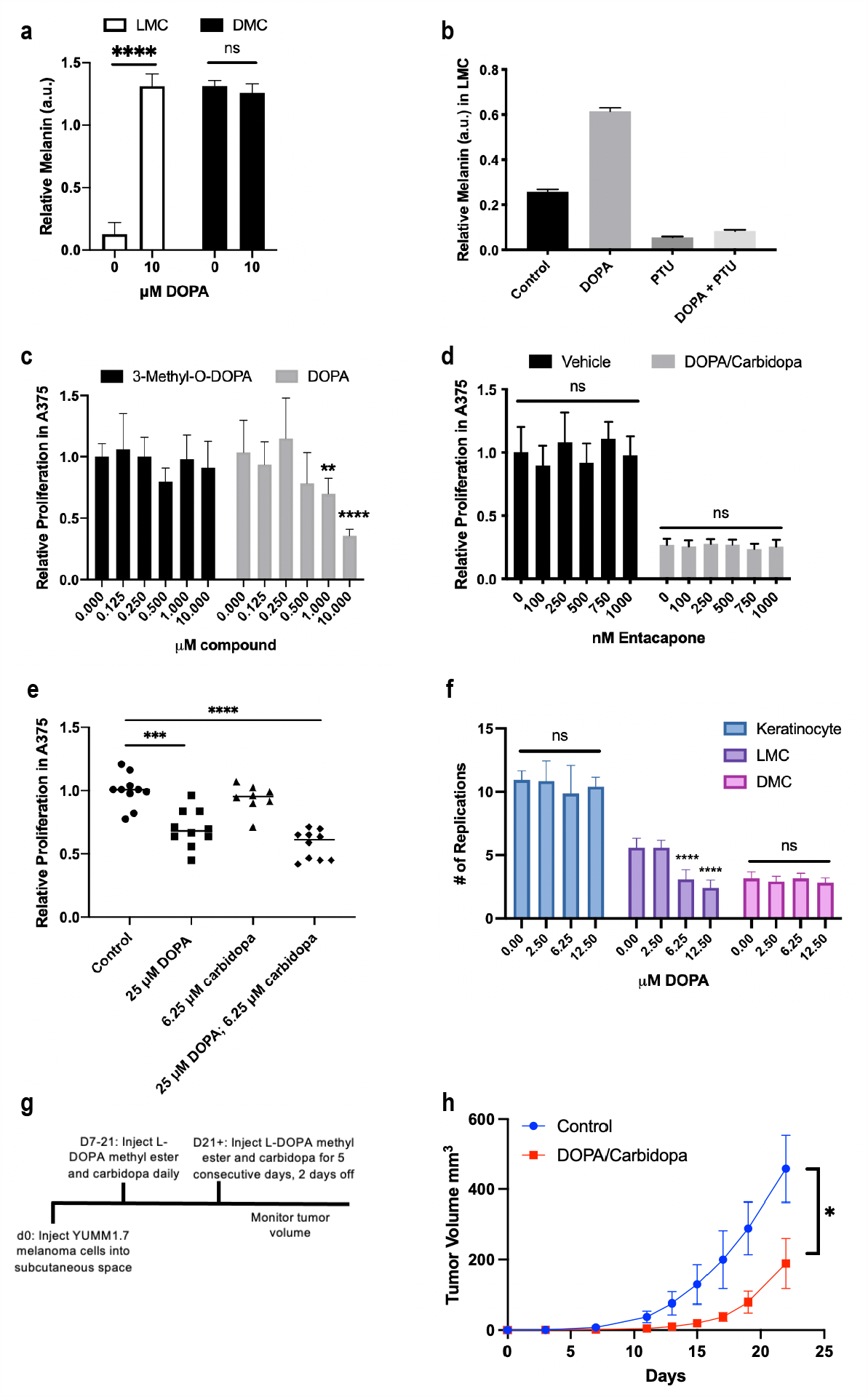
DOPA inhibits proliferation selectively in LMC, but not DMC. (a) Relative melanin content of lightly pigmented melanocytes (LMC) and darkly pigmented melanocytes (DMC) treated with 10 µM L-DOPA relative to cell number. P-value **** <0.0001, ns= not significant. n=3 (b) Melanin content of LMC treated with either 25 µM L-DOPA, 75 µM phenylthiourea (PTU), or both relative to cell number. n=3 (c) Proliferation of A375 human melanoma treated with a dose curve of 3-O-Methyl-DOPA or L-DOPA. P-value ** = 0.0017, **** <0.0001. n=3 (d) A375 treated with either vehicle or combination DOPA/carbidopa and an increasing concentration of entacapone, a catechol-O-methyltransferase (COMT) inhibitor to block conversion of L-DOPA to 3-O-Methyl-DOPA. n=3 (e) Proliferation of A375 human melanoma treated with 25 µM L-DOPA, 6.25 µM carbidopa, or a combination. P-value *** = 0.0001, **** <0.0001. n=5 (f) Proliferation of primary human keratinocytes and melanocytes treated with increasing concentrations of DOPA up to the saturating dose of12.5 µM. p-value **** <0.0001. n=3. (g) Experimental timeline of combination DOPA and carbidopa treatment of human melanoma cells, n=5 per group. (h) YUMM1.7 murine melanoma growth in SCID mice treated with vehicle or 300 mg/kg L-DOPA methyl ester and 75 mg/kg carbidopa. P-value * = 0.031. n=5 for each group.

**Supplemental Figure 3:**
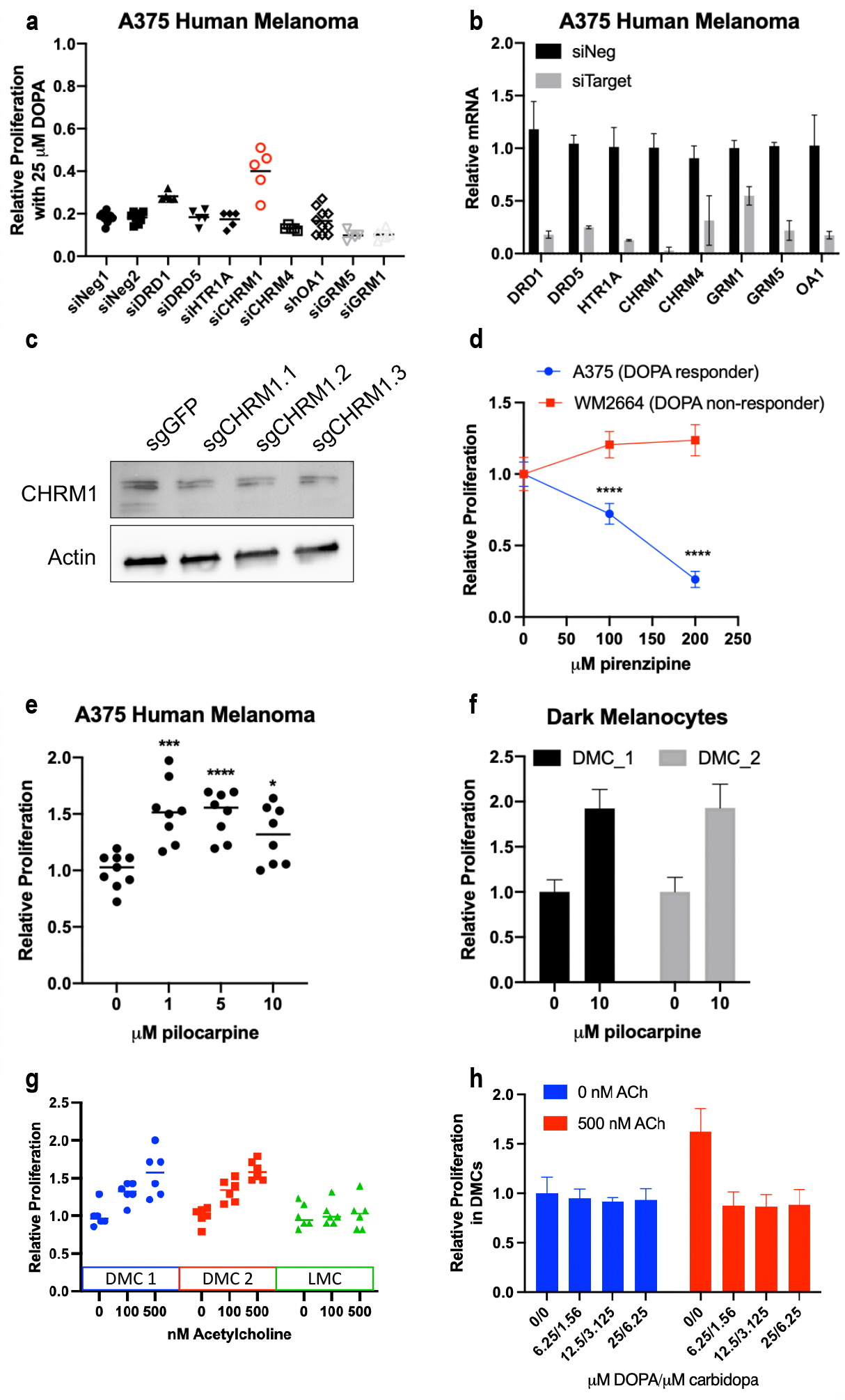
CHRM1 antagonism inhibits melanoma growth. (a) Pooled siRNA against top hits from PRESTO-Tango screen in A375 human melanoma in the presence of 25 µM L-DOPA and 6.25 µM carbidopa. Technical replicates, n=5. (b) qPCR from A375 treated with siRNA pools confirming gene knockdown. (c) Western blot for CHRM1 in A375 transduced with Cas9 and individual CHRM1 targeting gRNA. (d) Proliferation of A375 human melanoma (DOPA responder) and WM2664 human melanoma (DOPA non-responder) in presence of pirenzepine (CHRM1 antagonist). p-value < 0.0001, n=3. (e) Proliferation of A375 melanoma cells in presence of pilocarpine (CHRM1 agonist). p-value *** = 0.0007, **** < 0.0001, * = 0.0125. n=3 (f) Pilocarpine treatment in two biologically different darkly pigmented melanocytes (DMCs). (g) Five day proliferation of DMC and LMC with increasing concentrations of acetylcholine. (h) Five day proliferation of DMCs treated with an increasing concentration of DOPA/carbidopa in presence or absence of 500 nM acetylcholine.

**Supplemental Figure 4:**
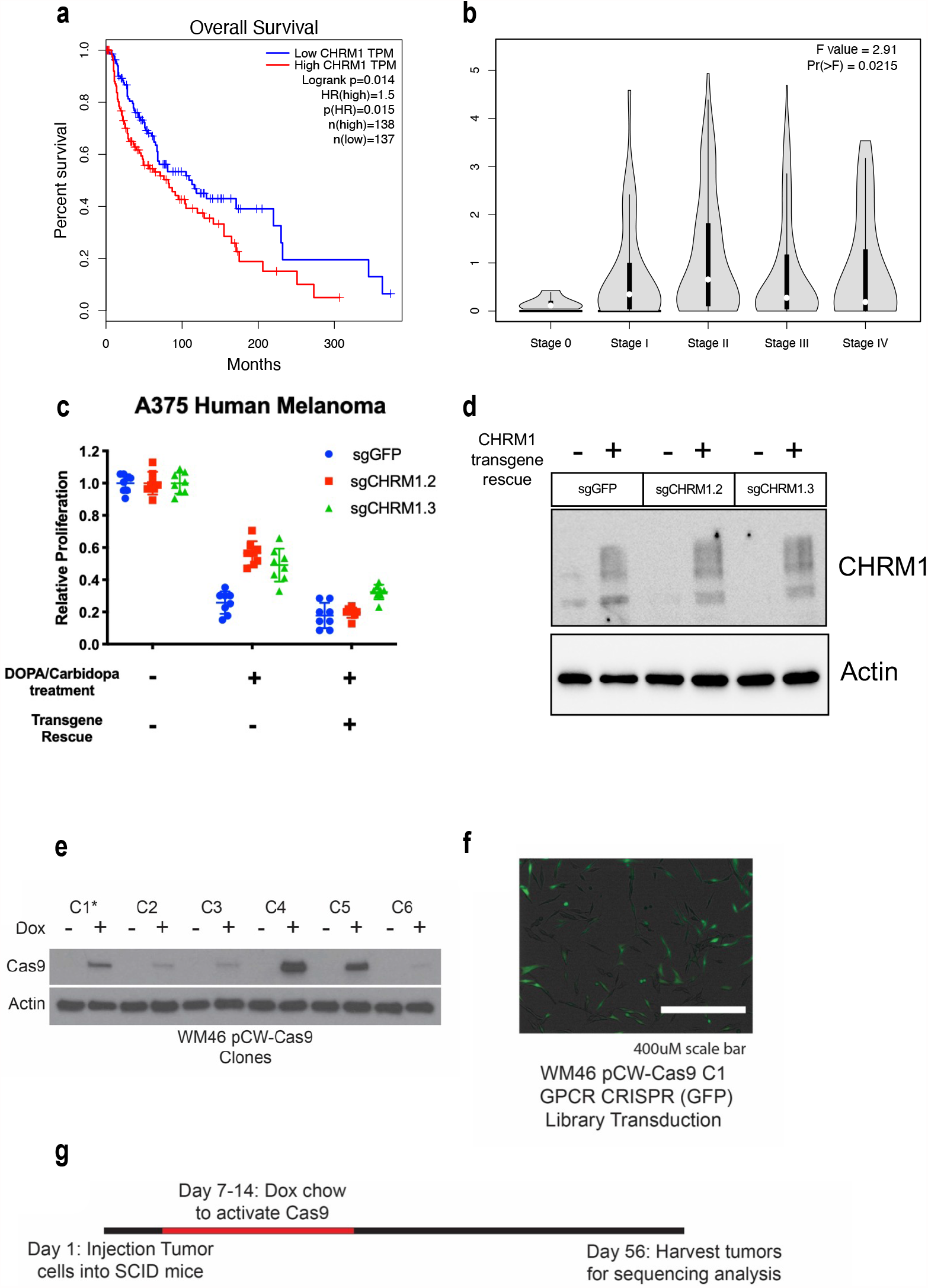
CHRM1 expression correlates with poor survival and invasiveness. (a) Kaplan-Meier overall survival in melanoma based on CHRM1 expression. Data obtained from with TCGA and GTEx datasets via GEPIA. (b) Pathological stage plot of CHRM1 in cutaneous melanoma via GEPIA. (c) Proliferation of A375 cells with CRISPR-Cas9 mediated CHRM1 depletion +/- CHRM1 transgene rescue treated with vehicle control or 25 µM L-DOPA and 6.25 µM carbidopa. n=5 (d) Western blot for CHRM1 from cells used in (c). (e) Western blot for doxycycline-inducible Cas9 protein in clonal populations of WM46 human melanoma cells to identify tightly controlled clones. C1 was picked for ***in vivo*** studies. (f) Fluorescence microscopy image of WM46 dCAS9 cells transduced with GPCR CRISPR library expressing GFP. (g) Experimental timeline of in vivo CRISPR screen.

**Supplemental Figure 5:**
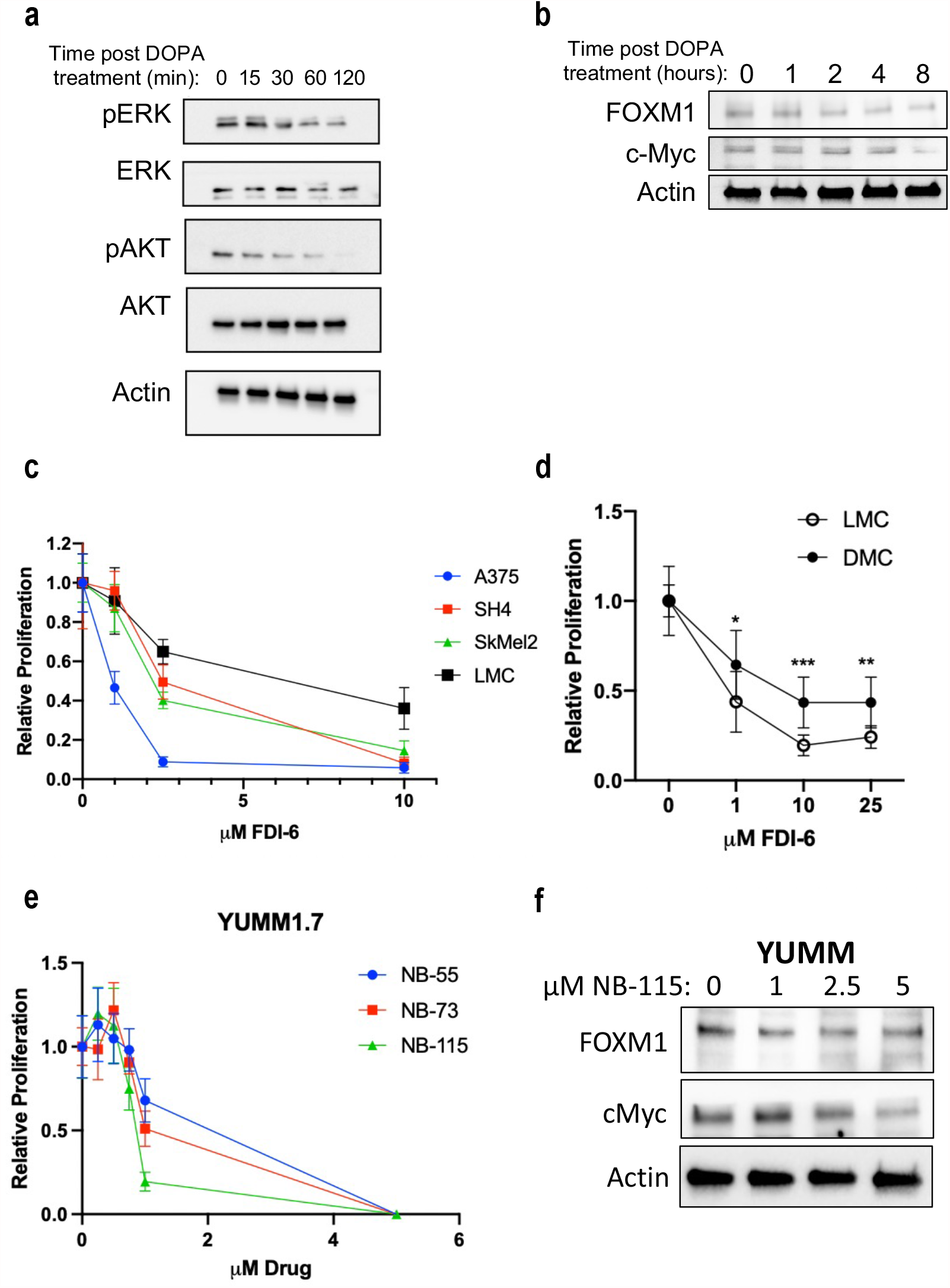
Pharmacologic FOXM1 antagonism inhibits MC and melanoma growth. (a) Western blot from A375 human melanoma treated with 25 µM L-DOPA and 6.25 µM carbidopa. (b) Western blot for FOXM1 and c-Myc in lysates from A375 human melanoma cells treated with 25 µM L-DOPA and 6.25 µM carbidopa. (c) Proliferation of human melanocytes and melanoma cells exposed to the FOXM1 inhibitor FDI-6, 4 day treatment. n= 3. (d) Relative proliferation of lightly pigmented melanocytes (LMC) and darkly pigmented melanocytes (DMC) after 4 days in presence of FDI-6. P-value * = 0.0164, *** = 0.0004, ** = 0.0025 by t-test. Technical replicates = 5, Biological replicate (both DMC and LMC) = 3. (e) Proliferation of YUMM1.7 melanoma in presence of new FOXM1 inhibitors, NB-55, NB-73 and NB-115. n = 5. (f) Western blot for FOXM1 and c-Myc in YUMM1.7 lysates after 24 hours of treatment with increasing NB-115 concentrations.

**Supplemental Table 1.**
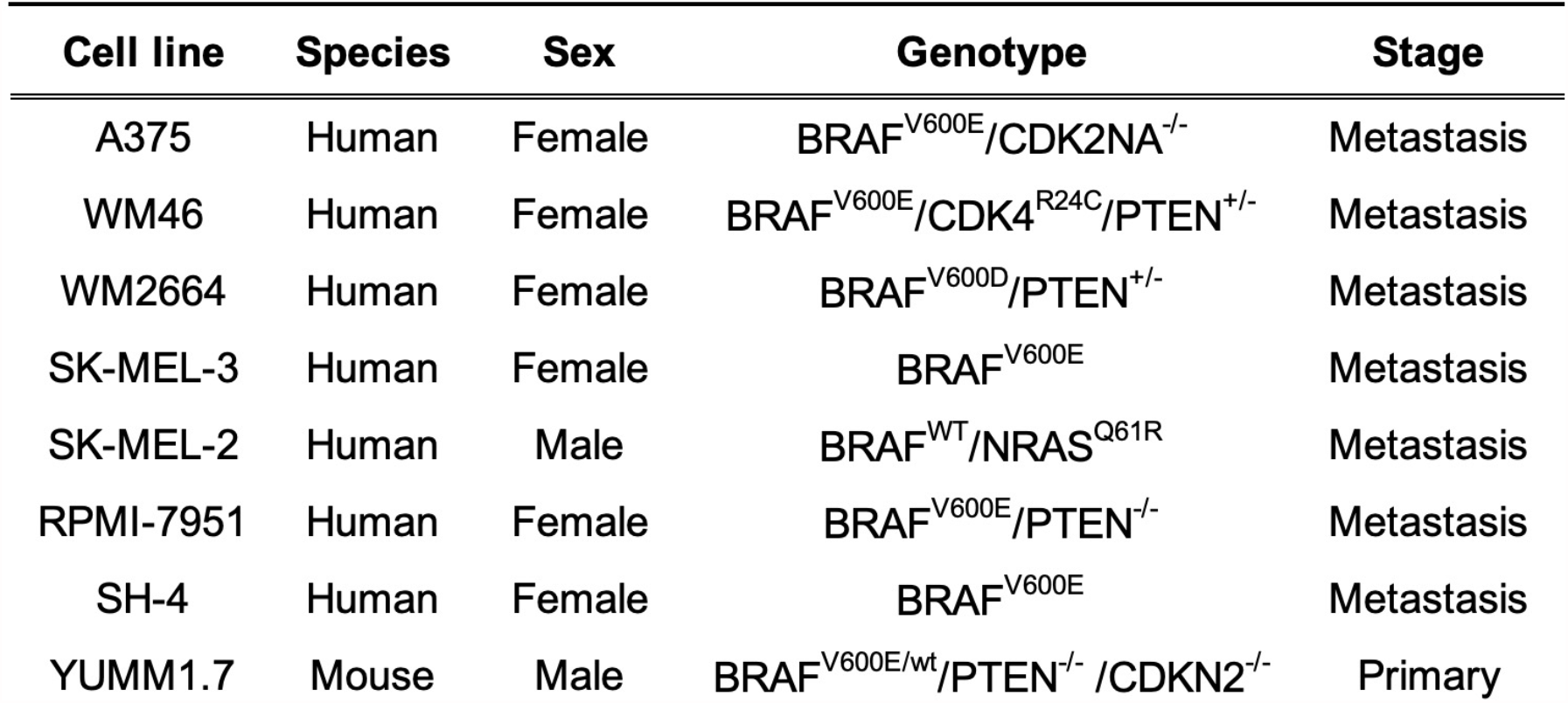
Profile of cell lines treated with combination DOPA/carbidopa.

**Supplemental Table 2.**
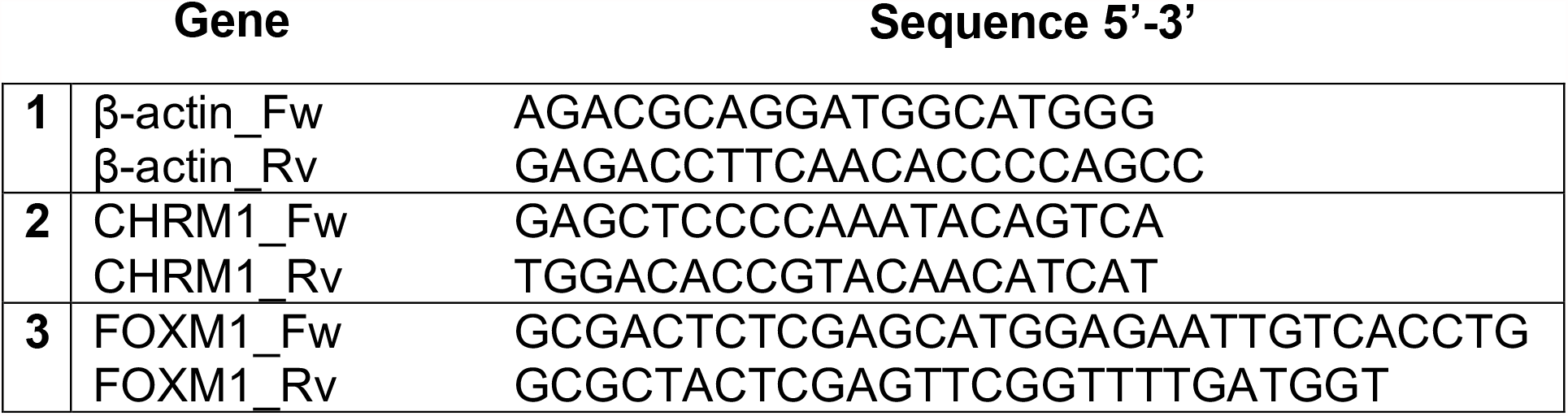
Primers used for Real-Time Quantitative PCRs.

