## Supplemental Figures for "CHRM1 is a Druggable Melanoma Target Whose Endogenous Activity is Determined by Inherited Genetic Variation in DOPA Production"

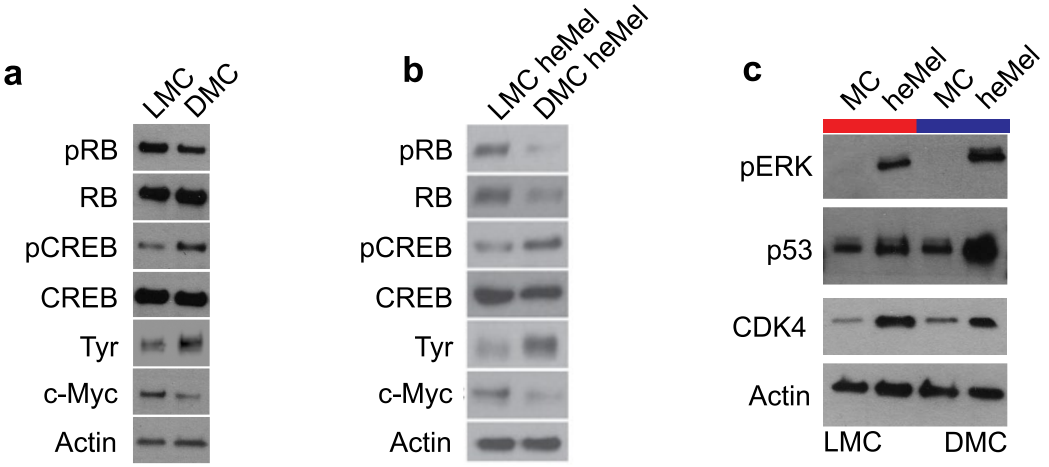


**Supplemental Figure 1: LMCs express lower levels of classical melanocyte differentiation markers, and higher levels of the proliferation driver and stem cell marker c-Myc than DMCs**. (a) Western blot of differentiation markers in representative lightly pigmented melanocytes (LMC) and darkly pigmented melanocytes (DMC). (b) Western blot of differentiation markers in representative lightly pigmented heMel (LMC heMel) and darkly pigmented heMel (DMC heMel). (c) Western blot of LMC and DMC transduced with heMel overexpression vectors.

Doepner_Supplemental_Fig_S2.

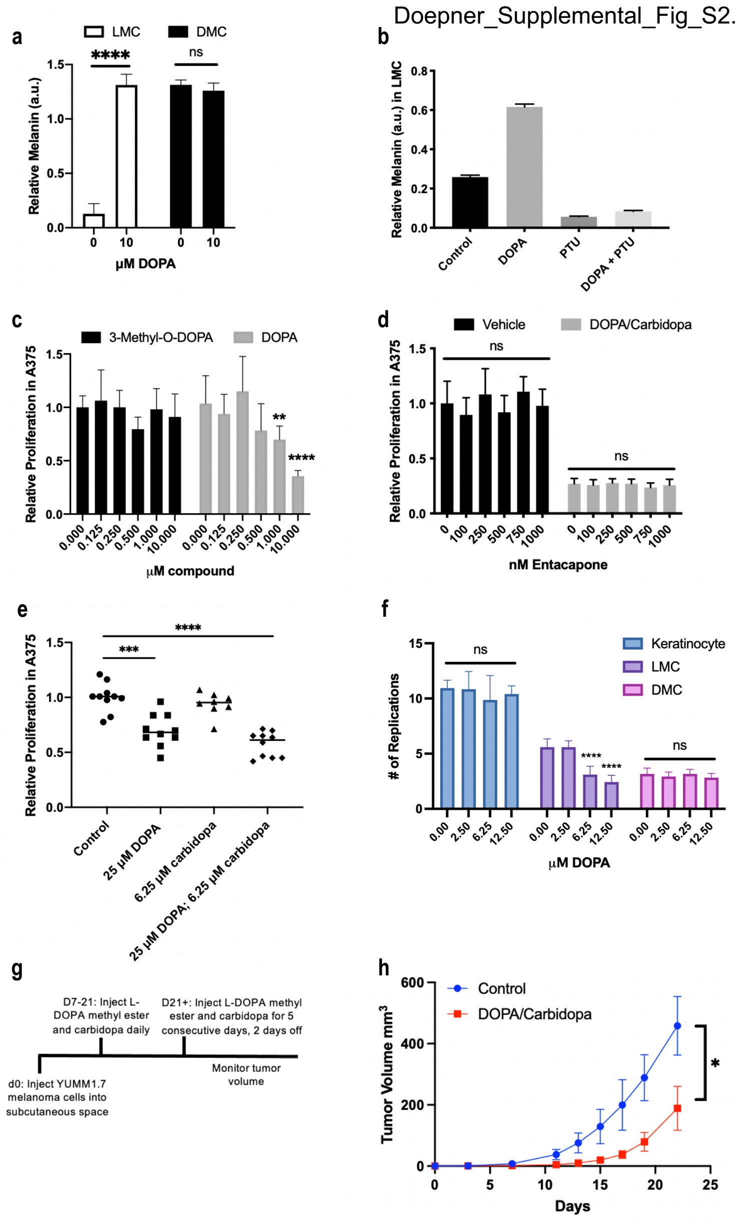


**Supplemental Figure 2: DOPA inhibits proliferation selectively in LMC, but not DMC.** (a) Relative melanin content of lightly pigmented melanocytes (LMC) and darkly pigmented melanocytes (DMC) treated with 10 µM L-DOPA relative to cell number. P-value **** <0.0001, ns= not significant. n=3 (b) Melanin content of LMC treated with either 25 µM L-DOPA, 75 µM phenylthiourea (PTU), or both relative to cell number. n=3 (c) Proliferation of A375 human melanoma treated with a dose curve of 3-O-Methyl-DOPA or L-DOPA. P-value ** = 0.0017, **** <0.0001. n=3 (d) A375 treated with either vehicle or combination DOPA/carbidopa and an increasing concentration of entacapone, a catechol-O-methyltransferase (COMT) inhibitor to block conversion of L-DOPA to 3-O-Methyl-DOPA. n=3 (e) Proliferation of A375 human melanoma treated with 25 µM L-DOPA, 6.25 µM carbidopa, or a combination. P-value *** = 0.0001, **** <0.0001. n=5 (f) Proliferation of primary human keratinocytes and melanocytes treated with increasing concentrations of DOPA up to the saturating dose of12.5 µM. p-value **** <0.0001. n=3. (g) Experimental timeline of combination DOPA and carbidopa treatment of human melanoma cells, n=5 per group. (h) YUMM1.7 murine melanoma growth in SCID mice treated with vehicle or 300 mg/kg L-DOPA methyl ester and 75 mg/kg carbidopa. P-value * = 0.031. n=5 for each group.

Doepner_Supplemental_Fig_S3.

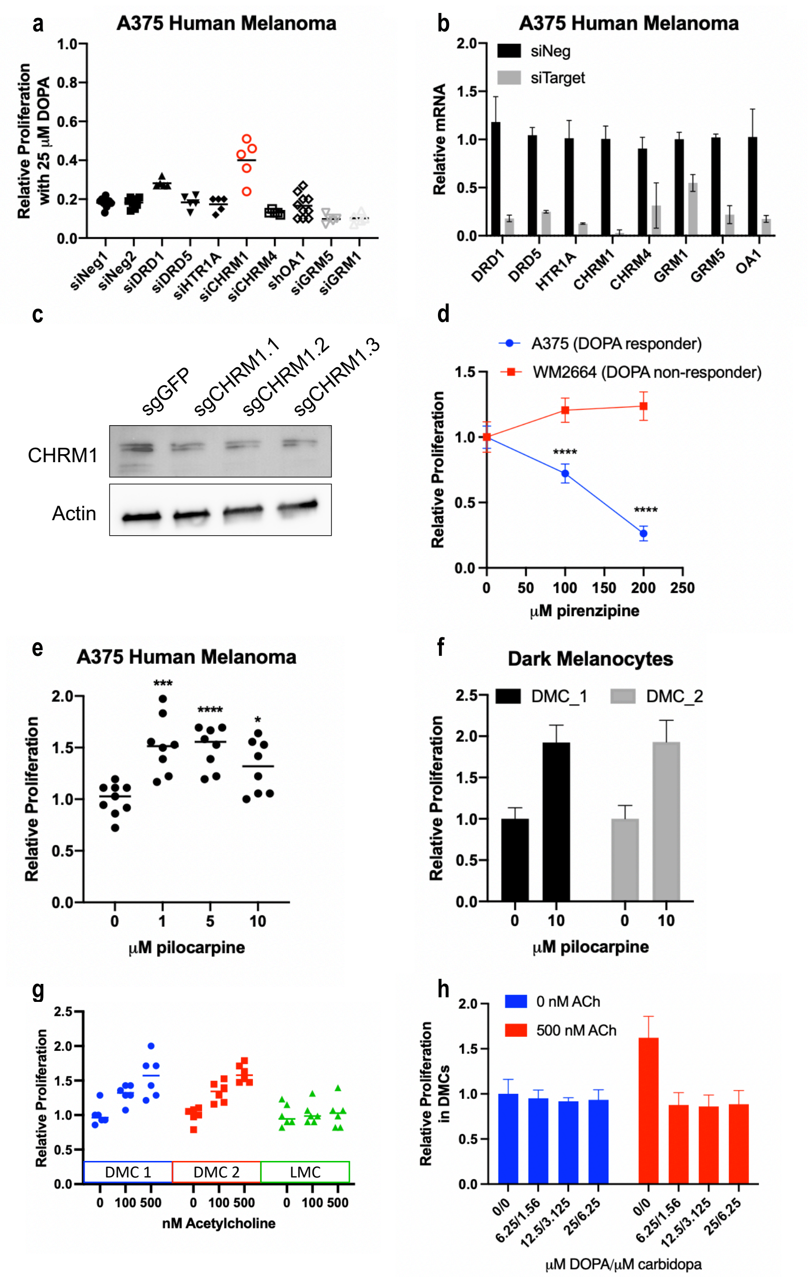


**Supplemental Figure 3: CHRM1 antagonism inhibits melanoma growth.** (a) Pooled siRNA against top hits from PRESTO-Tango screen in A375 human melanoma in the presence of 25 μM L-DOPA and 6.25 μM carbidopa. Technical replicates, n=5. (b) qPCR from A375 treated with siRNA pools confirming gene knockdown. (c) Western blot for CHRM1 in A375 transduced with Cas9 and individual CHRM1 targeting gRNA. (d) Proliferation of A375 human melanoma (DOPA responder) and WM2664 human melanoma (DOPA non-responder) in presence of pirenzepine (CHRM1 antagonist). p-value < 0.0001, n=3. (e) Proliferation of A375 melanoma cells in presence of pilocarpine (CHRM1 agonist). p-value *** = 0.0007, **** < 0.0001 , * = 0.0125. n=3 (f) Pilocarpine treatment in two biologically different darkly pigmented melanocytes (DMCs). (g) Five day proliferation of DMC and LMC with increasing concentrations of acetylcholine. (h) Five day proliferation of DMCs treated with an increasing concentration of DOPA/carbidopa in presence or absence of 500 nM acetylcholine.

Doepner_Supplemental_Fig_S4.

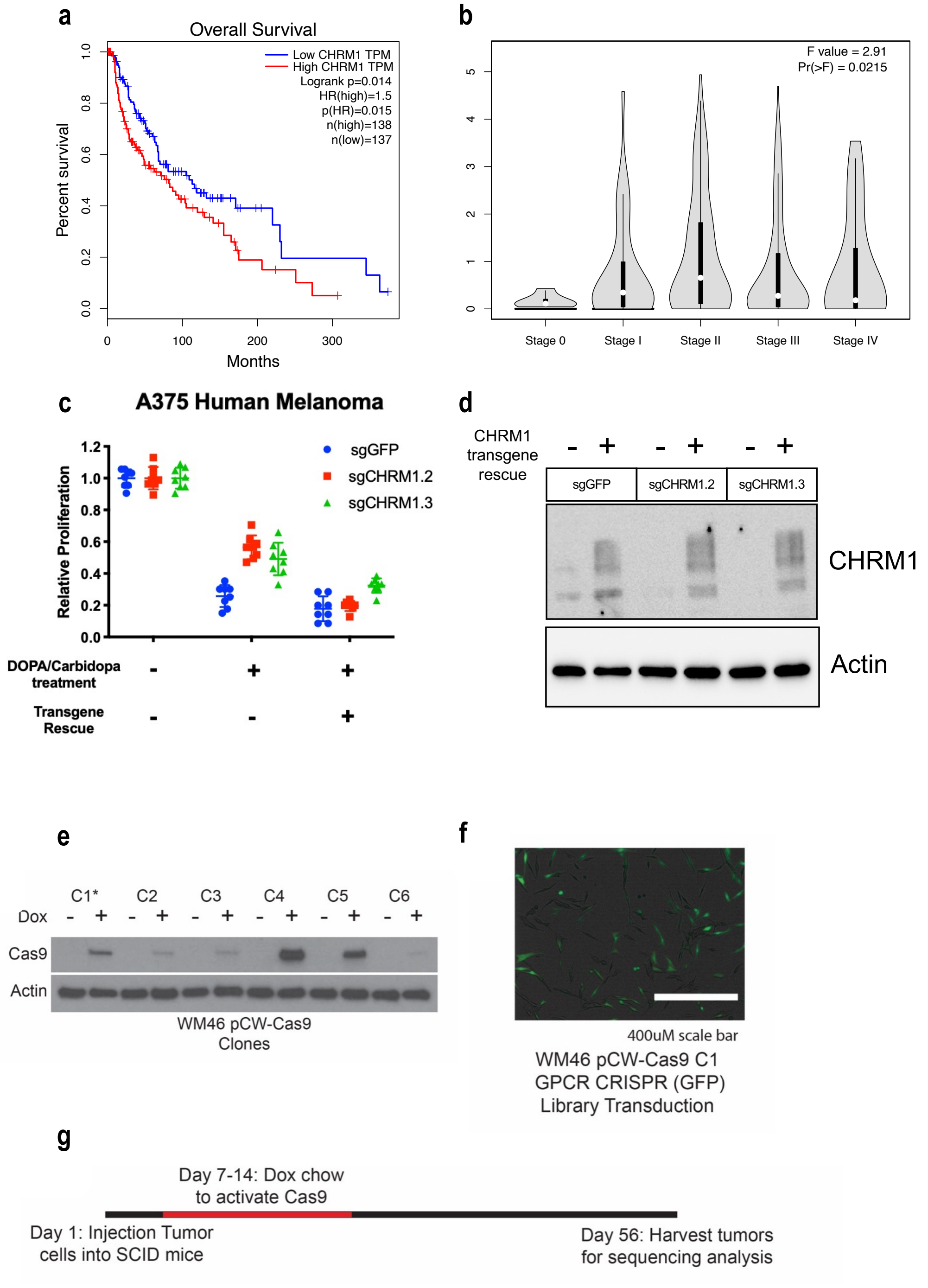


**Supplemental Figure 4: CHRM1 expression correlates with poor survival and invasiveness.** (a) Kaplan-Meier overall survival in melanoma based on CHRM1 expression. Data obtained from with TCGA and GTEx datasets via GEPIA. (b) Pathological stage plot of CHRM1 in cutaneous melanoma via GEPIA. (c) Proliferation of A375 cells with CRISPR-Cas9 mediated CHRM1 depletion +/- CHRM1 transgene rescue treated with vehicle control or 25 μM L-DOPA and 6.25 μM carbidopa. n=5 (d) Western blot for CHRM1 from cells used in (c). (e) Western blot for doxycycline-inducible Cas9 protein in clonal populations of WM46 human melanoma cells to identify tightly controlled clones. C1 was picked for ***in vivo*** studies. (f) Fluorescence microscopy image of WM46 dCAS9 cells transduced with GPCR CRISPR library expressing GFP. (g) Experimental timeline of in vivo CRISPR screen.

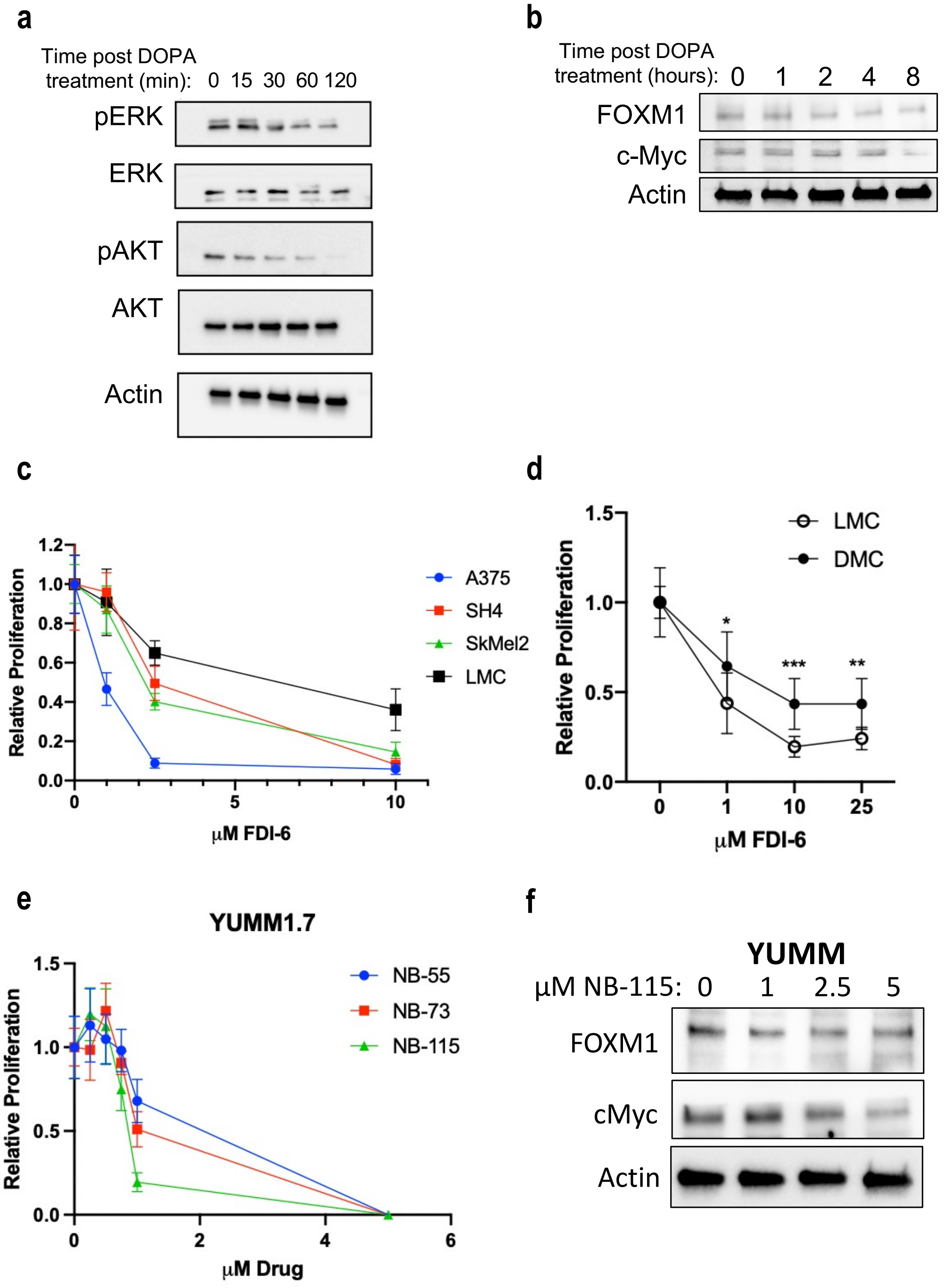


**Supplemental Figure 5: Pharmacologic FOXM1 antagonism inhibits MC and melanoma growth.** (a) Western blot from A375 human melanoma treated with 25 µM L-DOPA and 6.25 µM carbidopa. (b) Western blot for FOXM1 and c-Myc in lysates from A375 human melanoma cells treated with 25 µM L-DOPA and 6.25 µM carbidopa. (c) Proliferation of human melanocytes and melanoma cells exposed to the FOXM1 inhibitor FDI-6, 4 day treatment. n= 3. (d) Relative proliferation of lightly pigmented melanocytes (LMC) and darkly pigmented melanocytes (DMC) after 4 days in presence of FDI-6. P-value * = 0.0164, *** = 0.0004, ** = 0.0025 by t-test. Technical replicates = 5, Biological replicate (both DMC and LMC) = 3. (e) Proliferation of YUMM1.7 melanoma in presence of new FOXM1 inhibitors, NB-55, NB-73 and NB-115. n = 5. (f) Western blot for FOXM1 and c-Myc in YUMM1.7 lysates after 24 hours of treatment with increasing NB-115 concentrations.

Doepner_Supplemental_Table1.

| Cell line | Species | Sex | Genotype | Stage |
| --- | --- | --- | --- | --- |
| A375 | Human | Female | $BRAF^{V600E}/CDK2NA^{-/-}$ | Metastasis |
| WM46 | Human | Female | $BRAF^{V600E}/CDK4^{R24C}/PTEN^{+/-}$ | Metastasis |
| WM2664 | Human | Female | $BRAF^{V600D}/PTEN^{+/-}$ | Metastasis |
| SK-MEL-3 | Human | Female | $BRAF^{V600E}$ | Metastasis |
| SK-MEL-2 | Human | Male | $BRAF^{WT}/NRAS^{Q61R}$ | Metastasis |
| RPMI-7951 | Human | Female | $BRAF^{V600E}/PTEN^{-/-}$ | Metastasis |
| SH-4 | Human | Female | $BRAF^{V600E}$ | Metastasis |
| YUMM1.7 | Mouse | Male | $BRAF^{V600E/wt}/PTEN^{-/-}$ $/CDKN2^{-/-}$ | Primary |

**Supplemental Table 1.** Profile of cell lines treated with combination DOPA/carbidopa.

| | Gene | Sequence 5’-3’ |
|---|---|---|
| 1 | β-actin_Fw<br>β-actin_Rv | AGACGCAGGATGGCATGGG<br>GAGACCTTCAACACCCCAGCC |
| 2 | CHRM1_Fw<br>CHRM1_Rv | GAGCTCCCCAAATACAGTCA<br>TGGACACCGTACAACATCAT |
| 3 | FOXM1_Fw<br>FOXM1_Rv | GCGACTCTCGAGCATGGAGAATTGTCACCTG<br>GCGCTACTCGAGTTCGGTTTTGATGGT |

**Supplemental Table 2.** Primers used for Real-Time Quantitative PCRs.
